# Towards a unified model of naive T cell dynamics across the lifespan

**DOI:** 10.1101/2022.01.07.475400

**Authors:** Sanket Rane, Thea Hogan, Edward S. Lee, Benedict Seddon, Andrew J. Yates

## Abstract

Naive CD4 and CD8 T cells are cornerstones of adaptive immunity, but the dynamics of their establishment early in life and how their kinetics change as they mature following release from the thymus are poorly understood. Further, due to the diverse signals implicated in naive T cell survival, it has been a long-held and conceptually attractive view that they are sustained by active homeostatic control as thymic activity wanes. Here we employ multiple experimental systems to identify a unified model of naive CD4 and CD8 T cell population dynamics across the mouse lifespan. We infer that both subsets divide rarely and progressively increase their survival capacity with cell age. Strikingly, this simple model captures naive CD4 T cell dynamics throughout life. In contrast, we find that newly generated naive CD8 T cells are lost more rapidly during the first 3–4 weeks of life, likely due to increased recruitment into memory. We find no evidence for elevated division rates in neonates, or for feedback regulation of naive T cell numbers at any age. We show how confronting mathematical models with diverse datasets can reveal a quantitative and remarkably simple picture of naive T cell dynamics from birth into old age.

## Introduction

Lifelong and comprehensive adaptive immunity depends upon generating naive CD4 and CD8 T cell populations with diverse repertoires of T cell receptors (TCRs). These must be established rapidly from birth and then maintained throughout life. In mice, the number of circulating naive T cells grows from tens of thousands at birth to tens of millions in several weeks, peaking at around 2 months of age^1,2^ and waning thereafter. Quantifying the relative contributions of thymic influx, and of loss and self-renewal across the lifespan, processes that either boost or preserve diversity, will therefore help us understand at a mechanistic level how the TCR repertoire is generated and evolves as an individual ages.

In adult mice, naive T cells have a mean lifespan of several weeks but a mean interdivision time of several years^2,3^. This difference in timescales leads to the conclusion that, in mice, most naive T cells never divide and that their numbers are sustained largely by thymic export, which in adult mice contributes 1-2% of the peripheral pool size per day^1–5^. However, the dynamics of the naive pool may be radically different early in life, and it is unclear whether the rules that govern naive T cell dynamics in adults are the same in neonates. Indeed, there is considerable evidence that this is not the case. First, studies suggest that neonatal mice are lymphopenic, a state which, when artificially induced, supports the rapid expansion of newly-introduced T cells through a mechanism referred to as lymphopenia-induced proliferation (LIP)^6–10^, and T cells transferred to healthy neonatal mice undergo cell divisions not observed in adult recipients^11,12^. However, the early establishment of naive compartments is still heavily reliant upon thymic output, since depletion of thymocytes in 2 week-old mice drives a rapid and transient 50-70% reduction of peripheral CD4 and CD8 T cell numbers^13^.

Second, memory T cell compartments are rapidly established in neonatal mice, which derives from the activation of naive T cells. For instance, we have shown that the rate of generation of memory CD4 T cells is elevated early in life, at levels influenced by the antigenic content of the environment^14^. This result suggests that high *per capita* rates of activation upon first exposure to environmental stimuli may increase the apparent rate of loss of naive CD4 T cells in neonatal mice. One might expect a similar process to occur with naive CD8 T cells, with substantial numbers of so-called ‘virtual’ memory CD8 T cells generated from naive T cells in the periphery soon after birth^15^. Together, these observations suggest that the average residence times of naive T cells differ in neonates and adults.

Third, the post-thymic age of cells in neonates is inevitably more restricted than in adults. Following the dynamic period of their establishment, there is evidence that naive T cells do not die or self-renew at constant rates but continue to respond or adapt to the host environment^16^. Recent thymic emigrants (RTE) are both functionally distinct from mature T cells^17,18^, may be lost at a higher rate than mature naive T cells under healthy conditions^9,19–21^, and respond differently to mature naive cells under lymphopenia^9^. In the early weeks of life, all naive T cells are effectively RTE. Phenotypic markers of RTE are poorly defined, however, and so without a strict definition of ‘recent’ it is difficult to reach a consensus description of their kinetics. It may be more appropriate to view maturation as a continuum of states, and indeed the net loss rates (the balance of loss and self-renewal) of both naive CD4^22^ and CD8^22,23^ T cells in mice appear to fall smoothly with a cell’s post-thymic age, a process we have referred to as adaptation^22^. Such behaviour will lead to increasing heterogeneity in the kinetics of naive T cells over time, as the population’s age-distribution broadens, and may also contribute to skewing of the TCR repertoire, through a ‘first-in, last-out’ dynamic in which older naive T cells become progressively fitter than newer immigrants^3^. A conceptually similar model, in which naive T cells accrue fitness with their age through a sequence of stochastic mutation events, has been used to explain the reduced diversity of naive CD4 T cells in elderly humans^24^.

Taken together, these results indicate that host age, cell age, and cell numbers may all influence naive T cell dynamics to varying degrees. When dealing with cross-sectional observations of cell populations, these effects may be difficult to distinguish. For example, the progressive decrease in the population-average loss rate of naive T cells observed in thymectomised mice^2^ may not derive from reduced competition, as was suggested, but may also be explained by adaptation or selective effects^22^. It is also possible that elevated loss rates of naive T cells early in life may not be an effect of the neonatal environment *per se*, but just a consequence of the nascent naive T cell pool being comprised almost entirely of RTE with intrinsically shorter residence times than mature cells. These uncertainties invite the use of mathematical models to distinguish different descriptions of naive T cell population dynamics from birth into old age.

Here we combine model selection tools with data from multiple distinct experimental systems to investigate the rules governing naive T cell maintenance across the full life span of the mouse. We used an established bone marrow chimera system to specifically measure and model production, division and turnover of naive T cells in adult mice. We then used an out-of-sample prediction approach to test and refine these models in the settings of the establishment of the naive T cell compartments in neonates, and – using a unique Rag/Ki67 reporter mouse model – characterising the dynamics of RTE and mature naive T cells. We find that naive CD4 T cells appear to follow consistent rules of behaviour throughout the mouse lifespan, dividing very rarely and with a progressive increase in survival capacity with cell age, with no evidence for altered behaviour in neonates. Naive CD8 T cells behave similarly, but with an additional, increased rate of loss during the first few weeks of life that may reflect high levels of recruitment into early memory populations. These models are able to explain diverse observations and present a remarkably simple picture in which naive T cells appear to be passively maintained throughout life, with gradually extending lifespans that compensate in part from the decline in thymic output, and with no evidence for feedback regulation of cell numbers.

## Results

### Naive CD4 and CD8 T cells divide very rarely in adult mice and expected lifespans increase with cell age

Recent reports from our group and others^3,22,23,25^ show that the dynamics of naive CD4 and CD8 T cells in adult mice and humans depend on cell age, defined to be time since they (or their ancestor, if they have divided) were released from the thymus. All of these studies found that the net loss rate, which is the balance of their rate of loss through death or differentiation, and self-renewal through homeostatic division, decreases gradually with cell age for both subsets. It is unknown whether these adaptations with age modulate the processes that regulate their residence time, or their ability to self-renew. To address this question we used a well-established system that we have used to quantify lymphocyte dynamics at steady state in healthy mice^26^ but with the addition of detailed measurements of cell proliferation activity throughout. Briefly, hematopoietic stem cells (HSCs) in the bone marrow (BM) are partially and specifically depleted by optimized doses of the transplant conditioning drug busulfan, and reconstituted with T and B cell depleted BM from congenic donor mice. Chimerism rapidly stabilises among progenitors in the bone marrow^27^ and thymus^3^ and is maintained for the lifetime of the mouse (Figure 1A). The host’s peripheral lymphocyte populations are unperturbed by treatment, and as donor T cells develop they progressively replace host T cells in the periphery through natural turnover. This system allows us to estimate the rates of influx into different lymphocyte populations and the net loss rates of cells within them; identify subpopulations with different rates of turnover; and infer whether and how these dynamics vary with host and/or cell age^3,27,28^. Here, we generated a cohort of busulfan chimeric mice who underwent BMT between 7 and 25 weeks of age. At different times post-BMT, we enumerated host and donor-derived thymocyte subsets and peripheral naive T cells from spleen and lymph nodes (Figure 1B; see Figure S1 for the flow cytometric gating strategy). We began by normalising the chimerism (fraction donor) within naive CD4 and CD8 T cells to that of DP1 thymocytes to remove the effect of variation across mice in the stable level of bone-marrow chimerism. This normalised donor fraction (*f_d_*) will approach 1 within a population if it turns over completely – that is, if its donor:host composition equilibrates to that of its precursor. Saturation at *f_d_* < 1 implies incomplete replacement, which can occur either through waning influx from the precursor population, or if older (host) cells persist longer than new (donor) cells, on average, implying cell-age effects on turnover or self-renewal. Previously we observed incomplete replacement of both naive CD4 and CD8 T cells in adult busulfan chimeric mice^3^, and excluded the possibility that this shortfall derived from the natural involution of the thymus, leading us to infer that the net loss rates (the balance of loss and self-renewal through division) of both subsets increase with cell age^3,22^. For the present study we used concurrent measurements of Ki67, a nuclear protein that is expressed following entry into cell cycle and is detectable for approximately 3-4 days afterwards^28,29^, and stratified by host and donor cells (Figure 1B, lower panels). We reasoned that this new information would enable us to determine whether cell-age effects are manifest through survival or self-renewal.

**Figure 1:**
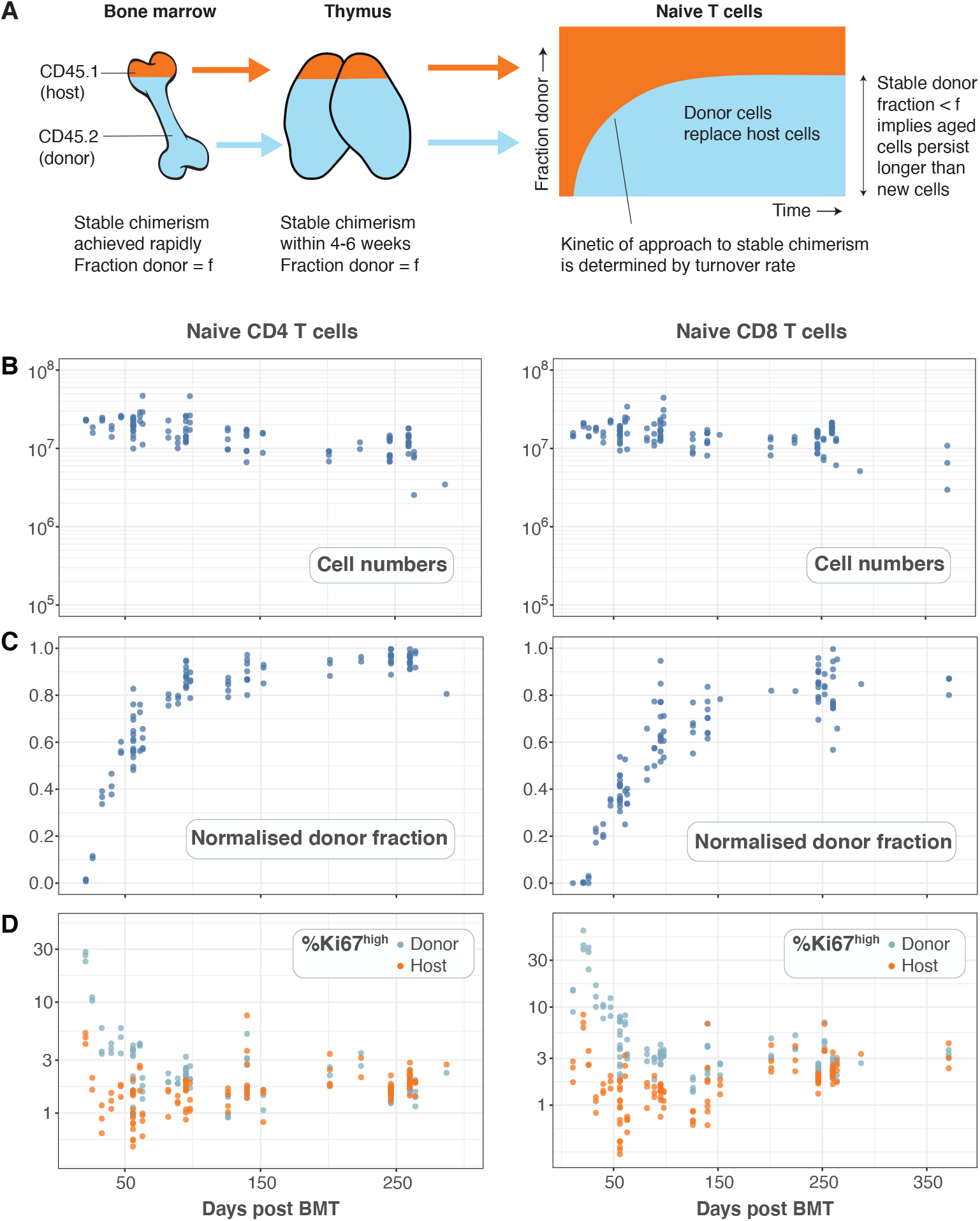
Dynamics of naive CD4 and CD8 T cells in busulfan chimeras. **(A)** The busulfan chimera system for studying lymphocyte population dynamics. **(B)** Observations of total naive T cell numbers derived from spleen and lymph nodes, their normalised donor fraction, and Ki67 expression among host and donor cells, in adult busulfan mice at various times post BMT.

To describe these data we explored variants of a structured population model in which either the rate of division or loss of naive T cells varies exponentially with their post-thymic age. These models are three dimensional linear partial differential equations (PDEs) that extend those we described previously^3,22^, allowing us to track the joint distribution of cell age and Ki67 expression within the population. We also considered a simpler variant of the age-structured models that explicitly distinguishes RTE from mature naive T cells, with a constant rate of maturation between two, and allowing each to have their own rates of division and loss^21,22^. We also considered models of homogeneous cell dynamics; the simplest ‘neutral’ model with uniform and constant rates of division and loss, and density-dependent models that allowed these rates to vary with population size. Models are illustrated schematically in Figure S2 and their formulations are detailed in Text S1. Each of them was fitted simultaneously to the measured time-courses of total naive CD4 or CD8 T cell numbers, the normalised donor fraction, and the proportions of donor and host cells expressing Ki67. To model influx from the thymus we used empirical functions fitted to the numbers and Ki67 expression levels of late stage single-positive CD4 and CD8 thymocytes (Text S2 and Figure S4). Since the rate of export of cells from the thymus is proportional to the number of single-positive thymocytes^19^, we used these functions to represent the rates of production of Ki67^hi^ and Ki67^low^ RTE with mouse age, up to a multiplicative constant which we estimated. The fitting procedure is outlined in Text S3, and detailed in ref. 27.

Our analysis confirmed support for the models of cell-age dependent kinetics, with all other candidates, including the RTE model, receiving substantially lower statistical support (Table S1). For naive CD4 T cells, we found strongest support for the age-dependent loss model (relative weight = 84%), which described their dynamics well (Figure 2A) and revealed that their daily loss rate declines as they age, halving roughly every 3 months (Table 1). For naive CD8 T cells the age-dependent division model was favoured statistically (relative weight = 87%; Figure 2B, dashed lines). However it yielded extremely low division rates, with recently exported cells having an estimated mean interdivision time of 18 months (95% CI: 14–25), and the division rate increasing only very slowly with cell age (doubling every 10 months). This model was therefore very similar to a neutral, homogeneous model and predicted that the normalised donor fraction approaches 1 in aged mice. This conclusion contradicts findings from our own and others’ studies that demonstrated that models assuming homogeneity in naive CD8 T cells failed to capture their dynamics in adult and aged mice (2-20 months old)^3,22,23^. A prediction of any progressive increase in cell division rates with age is also inconsistent with observations of the frequencies of T cell receptor excision circles (TRECs) in adult mice. TRECs are stable remnants of DNA generated during T cell receptor rearrangement, which are only diluted through mitosis; den Braber *et al*.^2^ found that the average TREC content of naive CD4 and CD8 T cells in mice aged between 12–125 weeks closely resembled the average TREC content of SP4 and SP8 thymocytes, suggesting that naive T cells in mice hardly divide. The statistical support for the age-dependent division model here may derive from the relatively sparse observations in aged mice in this dataset, which define the asymptotic replacement fraction. For the next phase of analysis we therefore retained the age-dependent loss model, which had the next highest level of support and was similar by visual inspection (Figure 2B, solid lines), as a candidate description of naive CD8 T cell dynamics.

**Figure 2:**
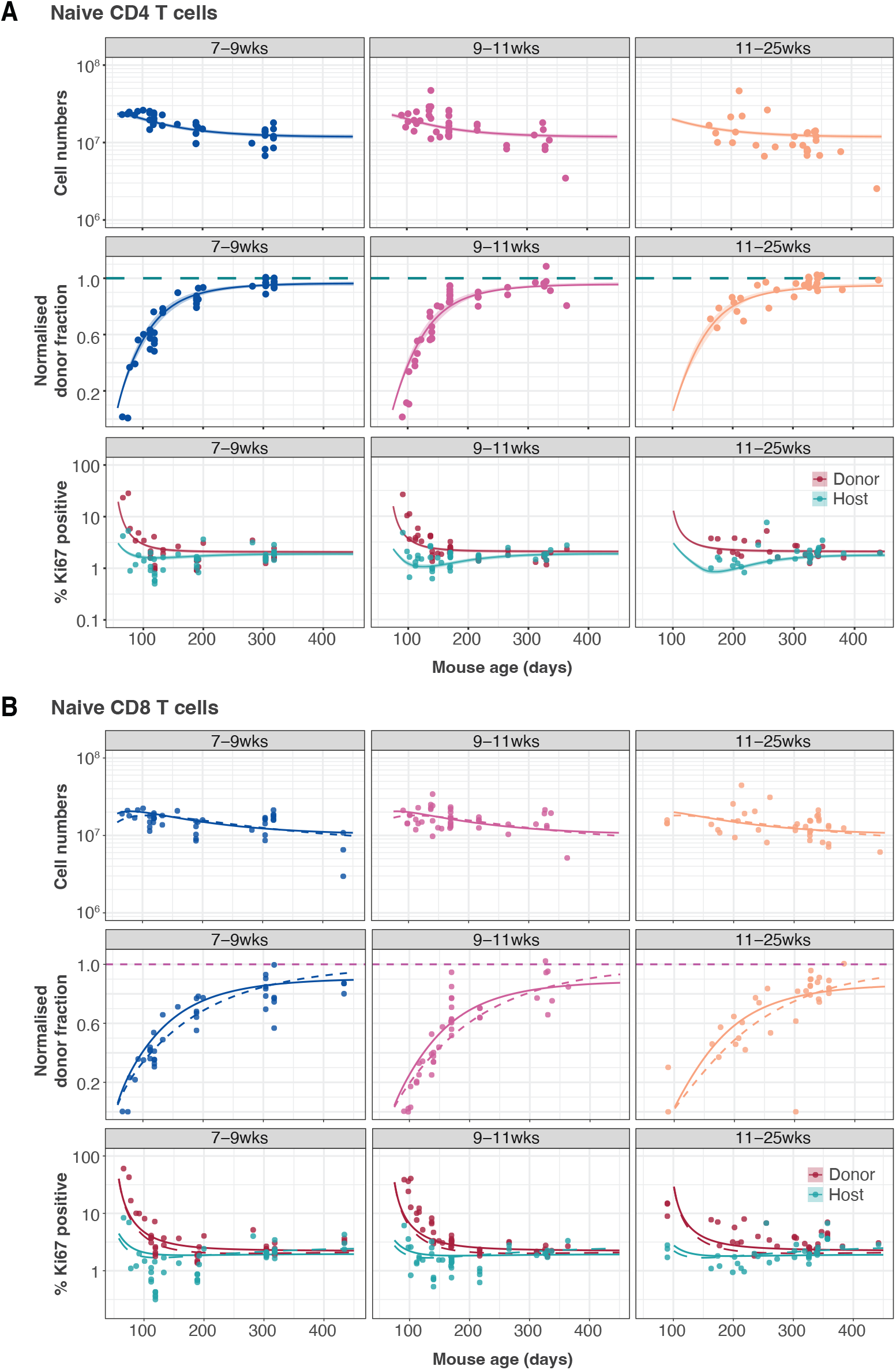
Modelling naive CD4 and CD8 T cell dynamics in adult busulfan chimeric mice. **(A)** The best fitting, age-dependent loss model of naive CD4 T cell dynamics describes the timecourses of their total numbers, chimerism and Ki67 expression in mice who underwent busulfan treatment and BMT at different ages (indicated in grey bars). **(B)** Naive CD8 T cell dynamics described by the age-dependent division model (dashed lines) and the age-dependent loss model (solid lines).

**Table 1:** Parameter estimates derived from fitting the age-dependent loss model to data from adult busulfan chimeric mice. Residence and interdivision times are defined as the inverses of the instantaneous loss rate (*δ*(*a*)) and the division rate (*ρ*), respectively. Posterior distributions of model parameters are shown in Figure S5.

| Population | Parameter | Estimate | 95% credible interval |
| --- | --- | --- | --- |
| Naive CD4 | Expected residence time of cells of age 0 (days) | 22 | 18 – 28 |
|  | Time taken for loss rate to halve (days) | 92 | 71 – 130 |
|  | Mean interdivision time (months) | 18 | 16 – 22 |
| Naive CD8 | Expected residence time of cells of age 0 (days) | 40 | 34 – 46 |
|  | Time taken for loss rate to halve (days) | 146 | 107 – 206 |
|  | Mean interdivision time (months) | 14 | 12 – 16 |

### Age-dependent loss models can describe RTE and mature naive CD4 and CD8 T cell kinetics in co-transfer experiments

To challenge these models further, we next confronted them with data from a study that compared the ability of RTE and mature naive (MN) CD4 and CD8 T cells to persist following co-transfer to an adult congenic recipient^9^. This study used a reporter mouse strain in which green fluorescent protein (GFP) expression is driven by *Rag2* gene expression elements, and is thus expressed throughout thymic development and for several days following export into the periphery. This is a long established and standard mouse model in which GFP expression is used as a surrogate marker of RTE status^30^. After transferring RTE and MN cells in equal numbers, the RTE:MN ratio within both CD4 and CD8 populations decreased progressively, falling by approximately 50% at 6 weeks (Figure 3), indicating that MN T cells persist significantly longer than RTE. We simulated this co-transfer using the models fitted to the data from the busulfan chimeric mice, and found that the age-dependent loss model predicted the trends in the CD4 and CD8 RTE:MN ratios (Figure 3A) while the age-dependent division model, which exhibited very weak age effects, predicted that the ratio would remain close to 1 (Figure 3B). We therefore favour models in which the expected residence times of naive CD4 and CD8 T cells increase with their age, while self-renewal occurs at very low, constant levels.

**Figure 3:**
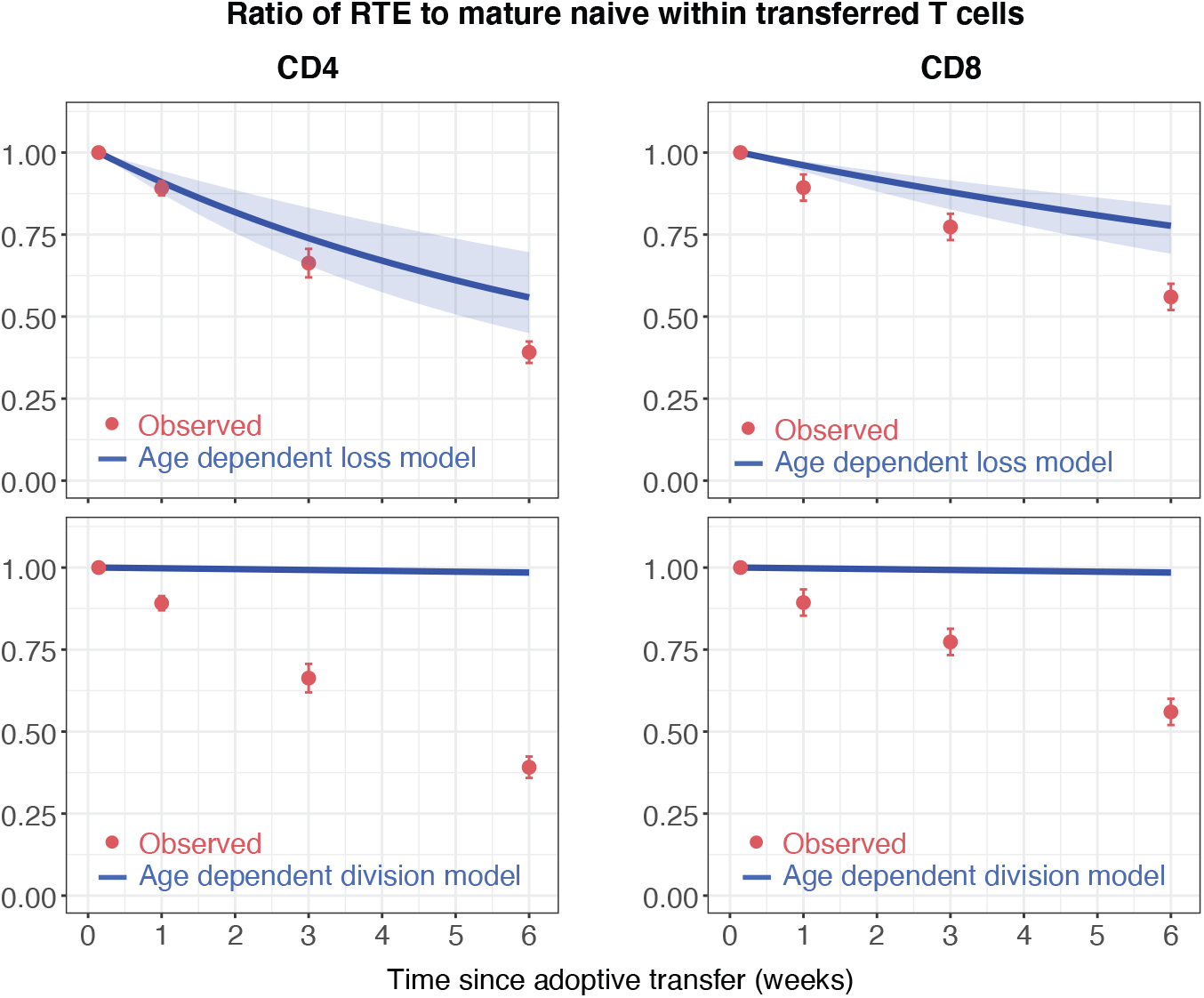
Models of age-dependent division fail to explain the survival kinetics of RTE and mature naive T cells. We simulated the co-transfer experiment described by Houston *et. al*.^9^ in which RTE from 5–9 week old Rag^GFP^ reporter mice were co-transferred with equal numbers of mature naive (MN) T cells from mice aged 14 weeks or greater to congenic recipients. Red points represent their observed RTE:MN ratios. The blue lines and envelopes shows the median and 5% and 95% quantiles of the ratio predicted by the model, sampling from the posterior distributions of parameters (shown in Figure S5) from the best-fitting age-dependent loss models (upper panels) and age-dependent division models (lower panels).

### Models parameterised using data from adult mice accurately predict the dynamics of naive CD4 T cells in neonates, but not of CD8 T cells

Next, we wanted to characterise the dynamics of naive CD4 and CD8 T cells during the first few weeks of life, and connect the two regimes to build unified models of the dynamics of these populations from birth into old age. Because it takes at least four weeks for peripheral donor-derived T cells to be detectable in busulfan chimeras, this system is not suitable for studying cell dynamics in young mice. Instead, we asked whether the models parameterised using data from adult mice could explain dynamics in young mice, and determine what (if any) modifications of the model were needed. We drew on two new data sets. One comprised the numbers and Ki67 expression of naive T cells derived from wild-type mice aged between 5 and 300 days. The other was derived from a cohort of Rag^GFP^ reporter mice, in which information about cell age can be gleaned from GFP expression levels. In this strain, intracellular staining for Ki67 is not possible without severely compromising GFP fluorescence. Therefore, we also introduced a Ki67^RFP^ reporter construct^31^ to the strain to generate Rag^GFP^Ki67^RFP^ dual reporter mice. Tracking GFP and RFP expression simultaneously therefore allows us to study the kinetics and division rates of RTE, which are enriched for GFP^+^ cells, and of mature naive T cells, which are expected to have largely lost GFP. We could then directly confront the models derived from adult mice with these new data.

Figure 4A and B show the numbers of naive CD4 and CD8 T cells and their Ki67 expression frequencies in three cohorts of mice – Rag^GFP^Ki67^RFP^ dual reporter mice aged between 10 and 120 days, WT mice, and adult busulfan chimeras in which host and donor cells were pooled. The red curves show the predictions of the cell-age-dependent loss models, which were fitted to the busulfan chimera data (red points) and extrapolated back to 1 day after birth. The dual reporter mice also yielded measurements of the co-expression of GFP and Ki67. To predict the kinetics of GFP^+^ Ki67^−^ and GFP^+^ Ki67^+^ proportions (Figure 4C and D) we needed to estimate only one additional parameter – the average duration of GFP expression. We assume that RTE become GFP-negative with first order kinetics at a rate defined both by the intrinsic rate of decay of GFP and the threshold of expression used to define GFP^+^ cells by flow cytometry. Our estimates of the mean duration of GFP expression within CD4 and CD8 RTE were similar (11 and 8 days, respectively). Details of our analysis are provided in Text S5.

**Figure 4:**
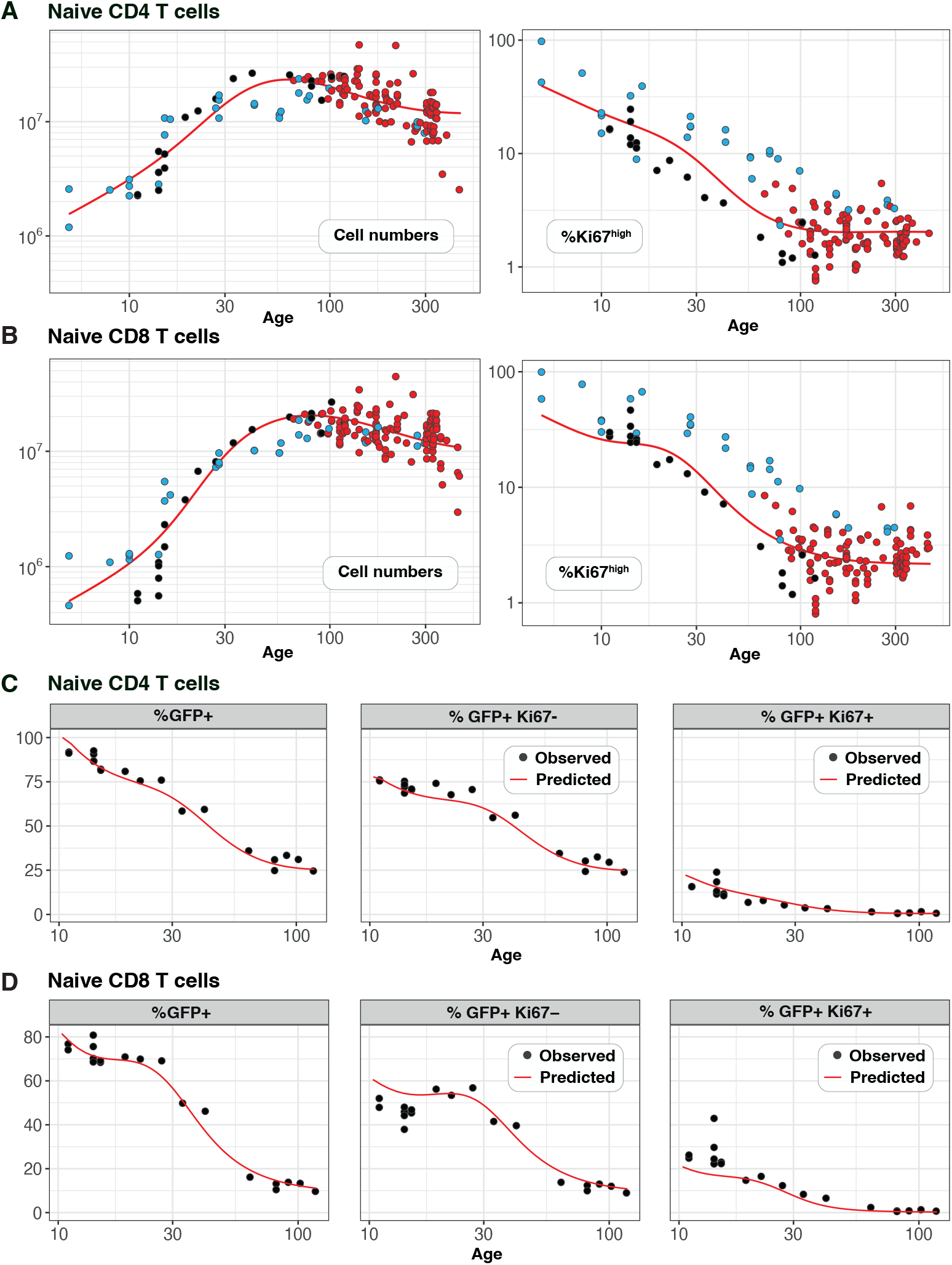
Predicting the kinetics of establishment of naive CD4 and CD8 T cell pools in early life. Panels **A and B**: For naive CD4 and CD8 cells, we extrapolated the age-dependent loss models (red curves) that were fitted to data from adult busulfan chimeric mice (red points) back to age 1 day. We compared these predicted trajectories with independent observations of naive T cell numbers and Ki67 expression in wild-type mice aged between 5-300 days (blue points), and from Rag^GFP^ Ki67^RFP^ reporter mice (black points). Panels **C and D**: We then estimated one additional parameter – the expected duration of GFP expression – by fitting the age-dependent loss model to the timecourses of total numbers of naive CD4 and CD8 GFP^+^ cells in these reporter mice (leftmost panels). We could then predict the timecourses of the percentages of GFP^+^ Ki67^+^ and GFP^+^ Ki67^−^ cells (centre and right panels).

Strikingly, the model of naive CD4 T cell dynamics in adult chimeric mice captured the total numbers and Ki67 expression of these cells in neonates remarkably well (Figure 4A), as well as the dynamics of Ki67^low^ and Ki67^hi^ RTE as defined by GFP expression (Figure 4C). This agreement indicates that the high level of Ki67 expression in naive CD4 T cells early in life does not reflect increased rates of division or LIP, but, is rather inherited from precursors within the neonatal thymus, a large fraction of which undergo cell division (Figure S4).

For naive CD8 T cells the cell-age-dependent loss model accurately predicted cell dynamics in both the reporter and wild-type mice back to approximately 3 weeks of age, but underestimated Ki67^hi^ frequencies in neonatal mice (Figure 4B, right panel), suggesting that naive CD8 T cells exhibit distinct dynamics very early in life. Intuitively this mismatch can be explained in two ways; either CD8 RTE in neonates are lost at a higher rate than in adults, or they divide more rapidly. In the former, a greater proportion of GFP^+^ Ki67^+^ RTE will be lost before they become Ki67^low^ and so the predicted proportion of cells that are GFP^+^ Ki67^low^ will be lower (Figure 4D, centre panel). In the latter, the proportion GFP^+^ Ki67^hi^ will increase (Figure 4D, right panel). Therefore, to explain naive CD8 T cell dynamics in neonates the basic model of cell-age-dependent loss in adults can be extended in two ways, modulating either the division or loss rate early in life.

### Naive CD8 T cells are lost at a higher rate in neonates than in adults

To distinguish between these possibilities we turned to a published study by Reynaldi *et al*.^23^, who used an elegant tamoxifen-driven CD4-Cre^ERT2^−RFP reporter mouse model to track cohorts of CD8 T cells released from the thymus into the peripheral circulation of animals of varying ages. In this model, a pulse of tamoxifen permanently induces RFP in cells expressing CD4, including CD4+CD8+ double positive thymocytes. The cohort of naive CD8 T cells deriving from these precursors continues to express RFP in the periphery and timecourses of their numbers in individual mice were estimated with serial sampling of blood^23^. These timecourses showed that the net loss rate of naive CD8 T cells appears to slow with their post-thymic age, and the rate of loss of cells immediately following release from the thymus appears to be greater in neonates than in adults (Figure 5A). Without measures of proliferation, these survival curves reflect only the net effect of survival and self-renewal. Nevertheless we reasoned that confronting our models with these additional data, and triangulating with inferences from other datasets, would allow us to identify a ‘universal’ model of naive CD8 T cell loss and division across the mouse lifespan.

**Figure 5:**
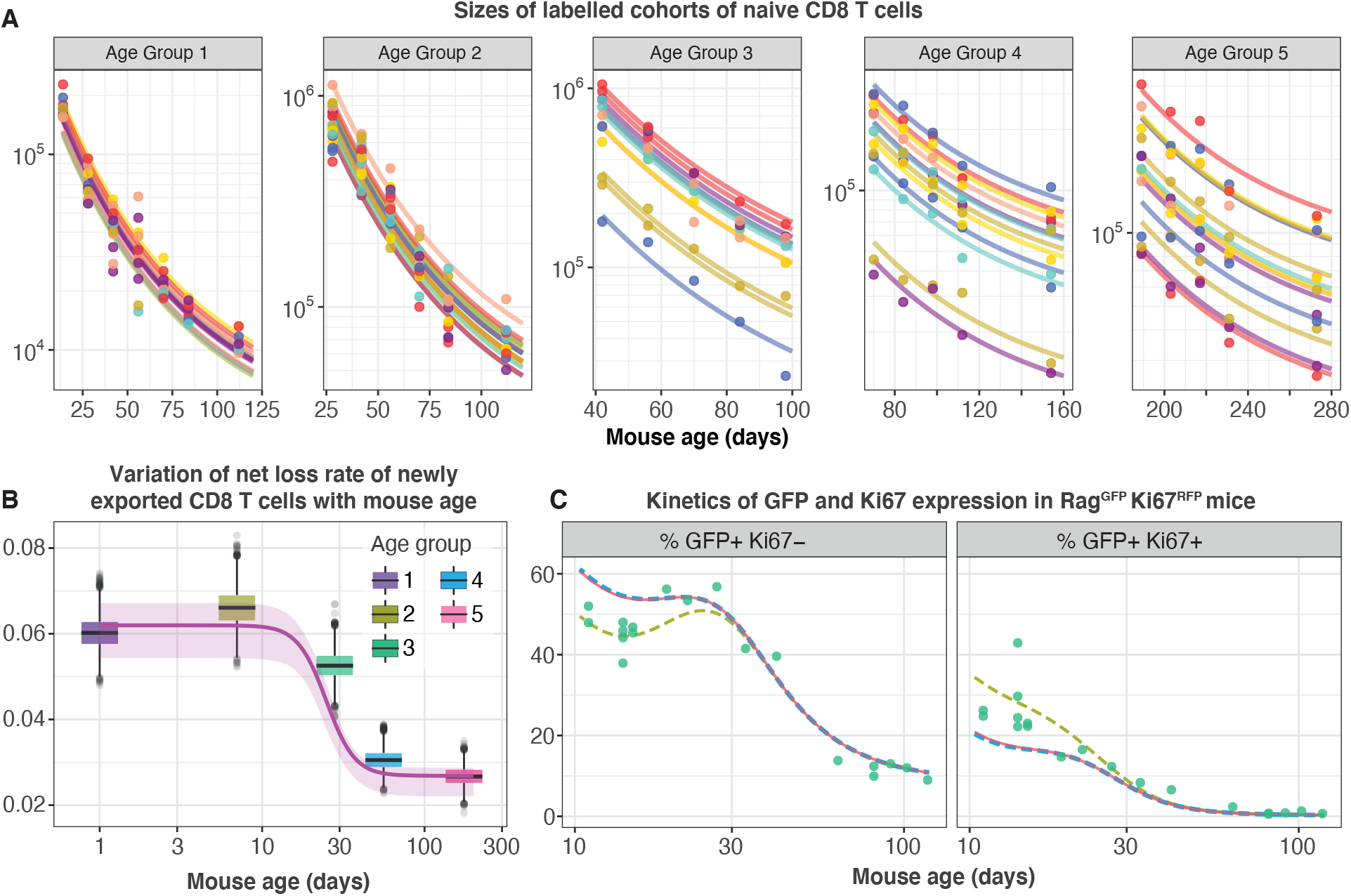
Tracking the persistence of cohorts of naive CD8 T cells *in vivo* – an analysis of data from Reynaldi *et al*.^23^. **(A)** Fitting the age-dependent loss model to the estimated numbers of time-stamped naive CD8 T cells in CD4-Cre^ERT2^ reporter mice treated with tamoxifen at different ages and sampled longitudinally. We used a hierarchical modelling framework and show mouse-specific fits to these timecourses (colours indicate different animals, dots are observations and lines are model fits). In the best fitting model, estimates of initial cell numbers were mouse-specific, while the net loss rate of cells of age 0 *i.e*. RTE (*λ*_0_ = *λ*(*a* = 0)) was specific to each mouse age group. **(B)** Corresponding estimates of *λ*_0_ for each age group of mice (black horizontal bars), with mouse-specific estimates (grey points) and the fitted, empirical description of *λ*_0_ with mouse age (see Text S6, eqn. S34). **(C)** Predicting the kinetics of the percentages of GFP^+^ Ki67^−^ and GFP^+^ Ki67^+^ CD8 T cells using the age-dependent loss model, including neonatal age effects in either the loss rate (green dashed line) or in the division rate (blue dashed line). The red line (partly concealed by the blue dashed line) shows the predictions of the original model fitted to the adult busulfan chimeric mice, with no mouse age effects.

We re-analysed the data from Reynaldi *et al*. using a Bayesian hierarchical approach (Text S6) to explain the variation in the kinetics of loss of these cohorts of cells across animals and age groups. Since there was no readout of cell division in this system, we simplified the cell-age-dependent loss model by combining division and loss into a net loss rate *λ*(*α*). We then fitted this model to the timecourses of labelled naive CD8 T cells across the different treatment groups. We tested four possibilities in which either the initial numbers of labelled cells (*N*_0_) and/or the net loss rate of cells of age 0(*λ*_0_) are varied across groups or animals as normally distributed hyper-parameters (Table S2). The model in which *N*_0_ was specific to each mouse and *λ*_0_ was specific to each age group gained 100% relative support (Table S2; fits in Figure 5A). This model confirmed that CD8 RTE are indeed lost at a significantly higher rate in the younger groups of mice (Figure 5B). We then described this decline in *λ*_0_ with mouse age empirically with a sigmoid (Hill) function, *λ*_0_(*t*) (Figure 5B, solid line) and used it to replace the discrete group-level variation in *λ*_0_ within the hierarchical age-structured model (Text S6). This ‘universal’ model, in which the loss rate of naive CD8 T cells declines with cell age but begins at higher baseline levels early in life, explained the data from Reynaldi *et al*. equally well (ΔLOO-IC < 6) and yielded visually indistinguishable fits.

This analysis shows that the baseline net loss rate of CD8 RTE declines from the age of ~3 weeks and stabilises at a level approximately 50% lower by age 9 weeks (Figure 5B). Therefore, newly exported naive CD8 T cells are at lost a higher rate in neonates than in adults, or they divide more slowly. Only the former is consistent with our inference from the Rag/Ki67 dual reporter mice. Indeed, we confirmed that simulating the age-dependent loss model from birth with a lower baseline division rate in neonates than in adults failed to improve the description of the early trajectories of the frequencies of GFP^+^ Ki67^−^ and GFP^+^ Ki67^−^ naive CD8 T cells (Figure 5C, blue dashed line). In contrast, increasing the baseline loss rate in neonates according to the function we derived from the data in Reynaldi *et al*. (Text S6) captured these dynamics well (Figure 5C, green dashed line).

In summary, we find that naive CD8 T cells rarely divide, increase their capacity to survive with cell age, and those generated within the first few weeks of life are lost at a higher baseline rate than those in adults.

### Ki67 expression within naive CD4 and CD8 T cells in adult mice is almost entirely a residual signal of intra-thymic proliferation

Our analyses are consistent with earlier reports that naive T cells in mice divide very rarely^2,3,32^. By explicitly modeling the kinetics of quiescent and recently-divided cells, we can also explain the apparently contradictory observation that more than 60% of naive CD4 and CD8 T cells express Ki67 early in life, declining to 2–3% by 3 months of age (Figure 4). We argue that this pattern, rather than being an indication of lymphopenia-induced proliferation early in life fading to low level but appreciable self-renewal in adults, is instead just a shadow of intrathymic division; Ki67 among peripheral naive T cells is almost entirely derived from cells that divided in the thymus and were exported within the previous few days. This conclusion emerged from the modelling of the busulfan chimera data but is also directly evident from the Rag2^EGFP^ Ki67^RFP^ reporter mice, in which Ki67-RFP expression among naive T cells was exclusively found on GFP^high^ peripheral RTE, and was a continuum of the expression by mature SP thymocytes (Figure 6A). This inheritance of expression from the thymus is also reflected in the high degree of correlation between the frequencies of Ki67-positive cells among SP thymocytes and peripheral naive T cells throughout life observed in WT mice (Figure 6B).

**Figure 6:**
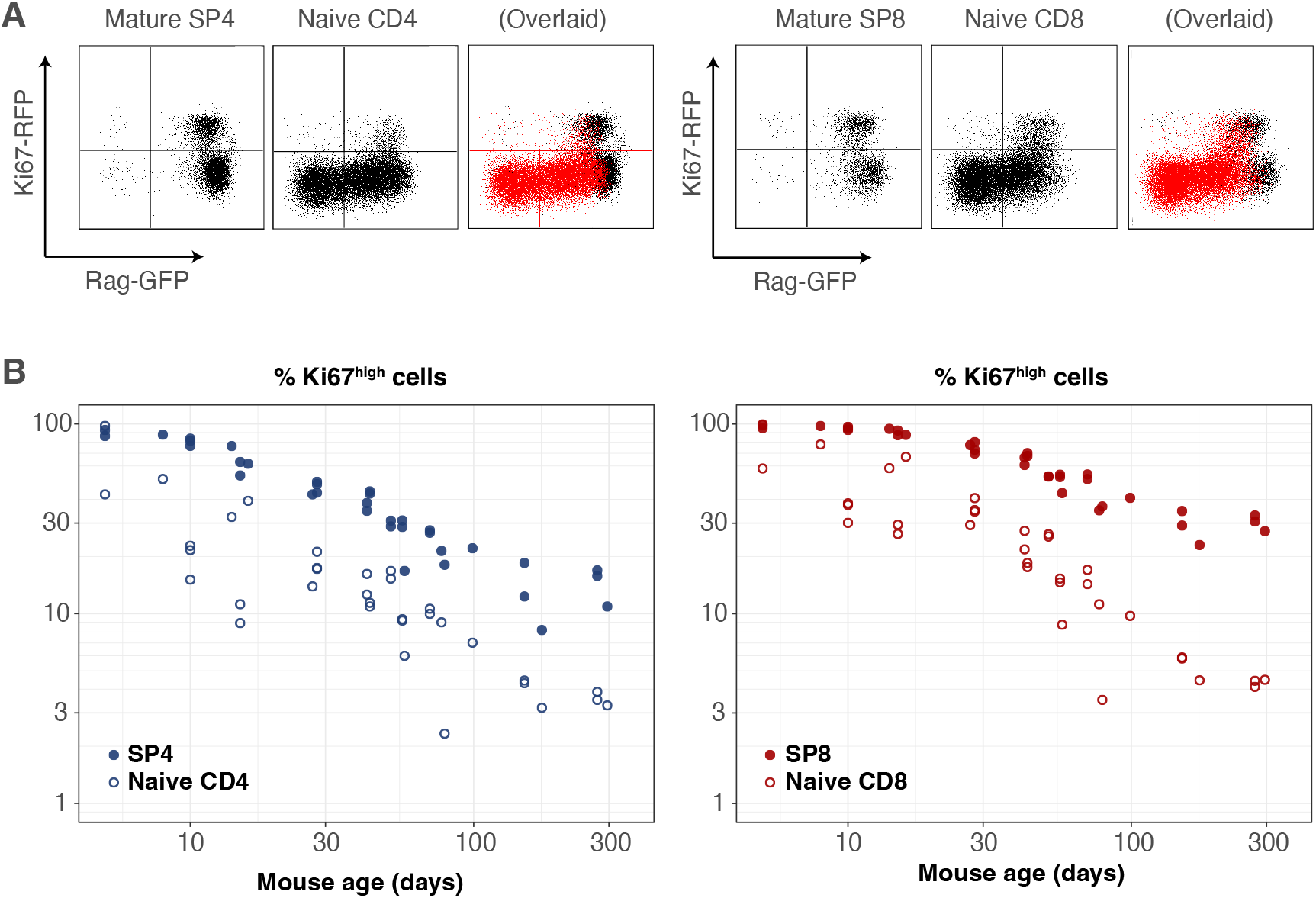
Markers of proliferation among naive T cells derive from very recent thymic emigrants. **(A)** Flow cytometry analyses of late stage single positive thymocytes and naive CD4 and CD8 T cells from lymph nodes in a 41 day-old Rag2^EGFP^ Ki67^RFP^ reporter mouse, showing that Ki67 expression among naive T cells is largely restricted to GFP^+^ RTE. In the ‘overlaid’ panels, naive T cells are shown in red and mature SP thymocytes in black. **(B)** Data from a cohort of wild-type mice showing that Ki67 levels in SP thymocytes and peripheral naive T cells correlate throughout life.

This result also gives an intuitive explanation of the trajectories of Ki67 expression within donor and host cells in the busulfan chimeric mice, which are distinct soon after BMT but converge after 6–12 months (Figure 1B and Figure 2). This behaviour does not derive from any intrinsic differences between host and donor T cells, but rather from the distinct age profiles of the two populations. Following BMT, the rate of production of host naive T cells declines substantially, as the procedure typically results in 80–90%replacement of host HSC with donor HSC. Since Ki67 is seen almost exclusively within very recent thymic emigrants, the frequency of Ki67-expressing host naive T cells then declines rapidly. Conversely, new donor-derived naive T cells are initially highly enriched for Ki67^hi^ cells. The frequencies of Ki67^hi^ cells within the two populations then gradually converge to pre-transplant levels as aged Ki67^low^ donor cells accumulate, and host-derived naive T cells equilibrate at lower numbers.

## Discussion

Our previous analyses suggested naive T cells operate autonomously and compensate cell-intrinsically for the gradual decline in thymic output with age by increasing their ability to persist with time since they leave the thymus in both adult mice^22^ and in humans^25^. Here we show through the modeling of a range of datasets that naive T cell adaptation in mice manifests primarily through a progressive decrease in their loss rate, and that they divide very rarely if at all, with mean interdivision times of at least 14 months. This means that throughout the mouse lifetime, newly made CD4 and CD8 RTE are lost at faster rates than their mature counterparts, predicting the preferential retention and accumulation of clones exported early in life. The lack of peripheral expansion combined with high levels of thymic export throughout life implies that the majority of the naive T cell repertoire is made up of small and long-lived TCR clones. This interpretation is consistent with studies showing enormous diversity within naive TCR repertoires in mice^33^ and supports the idea that any hierarchy within it is shaped by the generational frequencies of individual clones in the thymus, rather than by peripheral expansions^34^. In contrast, self-renewal of naive T cells is appreciable in humans and may contribute to the skewness of their TCR repertoires^35–37^. However, we observed a remarkably high degree of intrathymic proliferation in young mice, with close to 100% of late-stage CD62L^hi^ SP thymocytes expressing Ki67 in neonates, declining to approximately 20% over the first 3 months of life (Figures 6B and S4). Ki67 expression within these mature SP populations derives exclusively from cell division after TCR rearrangement and positive selection is complete. This observation implies that naive T clones generated in neonatal mice, which will ultimately be over-represented in older mice, may be substantially larger on average than those exported from adult thymi.

We showed agreement with the trends demonstrated by Houston *et al*.^9^, in which mature naive (MN) CD4 and CD8 T cell persist longer than RTE. However our simulations of their co-transfer experiment yielded somewhat lower levels of enrichment of MN cells (Figure 3A). This mismatch may derive in part from differences in the age-distributions of MN T cells in our simulations and their experiments. We considered cells of ages >21 days in 17 week old mice to be MN T cells. Houston *et al*. sourced MN T cells either from mice aged 14 weeks or older that were thymectomized at least 3 weeks prior to transfer, or from >12 week old Rag^GFP^ reporter mice. It is likely that their transferred MN cells were therefore enriched with older cells, giving a more pronounced disparity with RTE in survival.

A study by van Hoeven *et al*.^21^ also demonstrated that CD4 RTE are lost more rapidly than MN CD4 T cells^21^. They estimated that the loss rate of CD4 RTE was 0.063 day^−1^, translating to a residence time of 15 days (95% CI: 9–26), and is roughly 4 times shorter than the 66 day (52–90) residence time of MN CD4 T cells. Our results agree closely. We estimate that CD4 RTE (cells of age 0) have an expected residence time of 22 (18–28) days, doubling approximately every 3 months, such that in a 12 week old mouse, the mean residence time of MN CD4 T cells aged 21 days or greater is ~60 days. In contrast, van Hoeven *et al*. concluded that naive CD8 T cells are a kinetically homogeneous population with a mean residence time of 76 (42–83) days. With our favoured age-dependent loss model, we estimate that CD8 RTE initially have an expected residence time of 40 (18–28) days, doubling every ~5 months. However, our predicted average residence time of MN CD8 T cells (aged >21 days) in a 12 week old mouse was approximately 76 days, which agrees with their estimate. We included a similar RTE/MN model in our analysis (illustrated in Figure S2) and found that for CD8 T cells it received statistical support comparable to a neutral model of constant division and loss, in line with their analysis. Therefore, our different conclusions may stem in part from the specification of our models. It would be instructive to analyse the data from their thymic transplantation and heavy water labelling studies with the age-structured models we consider here.

The pioneering studies by Berzins *et al*. showed substantial and proportional increases in T cell numbers in mice transplanted with 2, 6 and 9 thymic lobes^19,20^. They concluded that this increase corresponds to the accumulation of RTE exported in the previous 3 weeks. In absence of any homeostatic regulation, the increase in the sizes of the naive CD4 and CD8 T cell pools under hyperthymic conditions is determined by the change in thymic output and by RTE lifespans. Our estimates of these lifespans (roughly 22 and 40 days for CD4 and CD8 respectively) are in line with their estimate of 3 weeks^20^. Indeed, simulating the transplantation of 6 thymic lobes using the age-dependent loss models and parameters derived from busulfan chimeric mice recapitulates their observations (Figure S6).

Reynaldi *et al*. used a novel fate-mapping system to demonstrate that the net loss rate of naive CD8 T cells (loss minus self-renewal) declines with their post-thymic age and is higher for CD8 RTE in neonates than in adults^23^. Our addition to this narrative was to reanalyse their data with a more mechanistic modeling approach to isolate the effects of division and loss, and to calculate a functional form for the dependence of the CD8 RTE loss rate on mouse age. In conjunction with our analysis of data from Rag/Ki67 dual reporter mice we inferred that the baseline loss rate naive CD8 T cells immediately following release from the thymus is higher in neonates than in adults, while the rate of division is close to zero throughout the mouse lifespan, and independent of host and cell age. The higher loss rate of CD8 RTE in neonates may derive from high rates of differentiation into memory phenotype cells rather than impaired survival. This idea is consistent with the rapid accumulation of virtual memory CD8 T cells in the periphery during the postnatal period^15^. However, we found no evidence for a similar process among naive CD4 T cells. While we recently showed that increasing the exposure to environmental antigens boosts the generation of memory CD4 T cell subsets early in life^14^, it may be that in the specific pathogen-free mice we studied here, any flux out of the naive CD4 T cell pool due to activation, which occurs before clonal expansion, was too low for our analysis to detect.

We do not explicitly model the mechanisms underlying adaptation in cell persistence. Modulation of sensitivity to IL-7 and signaling via Bcl-2 associated molecules have been implicated in increasing naive T cell longevity^9,38^ and is consistent with the outcome of co-transfer experiments. It is also possible that increased persistence derives additionally from a progressive or selective decrease in naive T cells’ ability to be triggered into effector or memory subsets. Studying the dynamics of naive T cells in busulfan chimeras generated using BM from TCR transgenic donor mice may help us untangle the contributions of survival and differentiation to the increase in their residence time with their age.

An alternative to the adaptation model is one of pure selection in which each cell’s survival capacity (loss rate) is determined during thymic development, drawn from a distribution, and subsequently fixed for its lifespan^22,39^. As an individual ages, naive T cells with intrinsically longer expected residence times will then be selected for. Indirect support for such a mechanism comes from the observation that low-level TCR signalling is essential for naive T cell survival^40,41^, suggesting that the ability to gain trophic signals from self-peptide MHC ligands may vary from clone to clone. We have also shown that different TCR-transgenic naive T cell clones have different capacities for proliferation in lymphopenic hosts^10^. However, one prediction of a selective model based on heterogeneity in TCR affinity alone is that TCR transgenic T cells co-transferred from young and old hosts would be lost at identical rates. One such experiment still saw that older cells exhibited a fitness advantage over younger ones^42^. Therefore, while we cannot rule out a pre-programmed (and possibly TCR-specific) element to each naive T cell’s life expectancy, it is clear that they also undergo progressive changes in their fitness, expressed in the adaptation models we have considered here.

Naive T cells proliferate under severely lymphopenic conditions in mice^7,8,10^ but acquire a memory-like phenotype^10,11,43,44^. There is some evidence that this process occurs in healthy neonatal mice^11^, suggesting that they are lymphopenic to some degree. However it may not contribute substantially to the production or maintenance of ‘truly’ naive T cells. In line with this conclusion, our analyses of neonatal mice revealed no evidence of increased rates of self-renewal, nor any reduction of cell loss rates, that would act to boost or preserve naive T cell numbers in the early weeks of life; neither did we need to invoke feedback regulation of their kinetics in adult mice. We showed previously that apparent density-dependent effects on naive T cell survival following thymectomy can also be explained by adaptive or selective processes^2,22^. Similarly, in humans, any regulation of a natural set-point appears to be incomplete at best; naive T cell numbers in HIV-infected adults typically do not normalise following antiretroviral therapy^45^, and recovery from autologous haematopoietic stem cell transplant results in persistent perturbations of T cell dynamics^46^. Dutilh *et al*.^47^ showed that explaining the kinetics of decline in TREC frequencies in human naive T cells requires an increase in either cell division or survival with age, as naive T cell numbers decline. They ascribed this to a density-dependent, homeostatic mechanism, but again cell-intrinsic adaptation or selection could underlie the phenomenon. Therefore, we speculate that the idea of naive T cell homeostasis over the life course, in the sense of compensatory or quorum sensing behaviour, may well be largely a theoretical concept. Selection pressures that shaped evolution of lymphocyte development are most likely to have been exerted on the establishment of T cell compartments and immunity that would support host survival to reproductive age, and would have little traction upon T cell behaviour into old age. Perhaps a better model, in both mice and humans, is the traditional understanding in which the thymus drives the generation of the bulk of the naive T cell pool in the early life, and thereafter naive T cell repertoires coast out into old age in a cell-autonomous manner.

## Materials and methods

### Generating busulfan chimeric mice

SJL.C57Bl6/J (CD45.1.B6) mice were treated with optimised low doses of busulfan to deplete HSC but leave peripheral T cell subsets intact. HSC were reconstituted with congenically-distinct, T-cell depleted bone marrow from C57Bl6/J donors to generate stable chimeras (Fig. 1A). Details of the protocols are given in ref. 26.

### Mice

Mki67tm1.1Cle/J (Ki67-RFP) mice were generously provided by the laboratory of Prof. Hans Clevers (Hubrecht Institute, KNAW and University Medical Centre Utrecht, Utrecht, The Netherlands). FVB-Tg(Rag2-EGFP) 1Mnz/J mice were from Jax Laboratories (strain 005688). Ki67-RFP x Rag2-EGFP F1 mice were subsequently backcrossed to a C57Bl6/J background for seven generations. Busulfan chimeric mice and wild-type control mice were housed in conventional animal facilities at the UCL Royal Free Campus, London, UK (UCL). Mice were housed in individually ventilated cages and drank irradiated water. Mouse experiments were subject to local ethical approval and under HO project license.

### Flow cytometry

Single cell suspensions were prepared from the thymus, spleen and lymph nodes of busulfan chimeric mice, wildtype control mice, or germ free mice. Cells were stained with the following monoclonal antibodies and cell dyes: CD45.1 FITC, CD45.2 FITC, CD45.2 AlexaFluor700, TCR-β APC, CD4^+^ PerCP-eFluor710, CD44 APC-eFluor780, CD25 PE, CD25 eFluor450, CD25 PE-Cy7, CD62L eFluor450, NK1.1 PE-Cy7 (all eBioscience), CD45.1 BV650, CD45.2 PE-Dazzle, TCR-β PerCP-Cy5.5 CD4^+^ BV711, CD44 BV785, CD25 BV650 (all Biolegend), CD62L BUV737 (BD Biosciences), LIVE/DEAD nearIR and LIVE/DEAD blue viability dyes. For Ki67 staining, cells were fixed using the eBioscience Foxp3/ Transcription Factor Staining Buffer Set and stained with either anti-mouse Ki67 FITC or PE (both eBioscience). Cells were acquired on a BD LSR-Fortessa flow cytometer and analysed with Flowjo software (Treestar). See Figure S1 for the gating strategy used to identify mature single positive thymocytes and peripheral naive subsets, and gates to measure Ki67 frequencies.

#### Mathematical modelling and statistical analysis

Using the data from adult busulfan chimeric mice, together with empirical descriptions of the pool sizes and Ki67^hi^ fraction within SP thymocytes to define thymic influx (Text S2), we fitted a set of candidate mathematical models (detailed in Text S1) to the time courses of total cell counts, normalised donor fraction and the fraction of cells that were Ki67^hi^ within donor and host subsets of naive CD4 and CD8 T cells. We used a Bayesian estimation approach using *R* and *Stan*. Code and data used to perform model fitting, and details of the prior distributions for parameters, are available at this linked [Github repository](https://github.com/sanketrane/T_cell_dynamics_birth-death). Models were ranked based on information criteria estimated using the Leave-One-Out (LOO) cross validation method^48,49^ (Text S3).

To predict neonatal dynamics of naive T cells, the models fitted to data from adult mice were extrapolated back to near birth (Text S4) and we constructed a mapping between cell age and GFP expression to predict the kinetics of GFP+ and Ki67+ cells in Rag^GFP^Ki67^RFP^ reporter mice aged between 11 days and 4 months (Text S5). To re-analyze longitudinal data from Reynaldi *et al*.^23^, tracking the survival of cohorts of naive CD8 T cells within different age groups of mice, we used a hierarchical Bayesian modelling approach (Text S6).

## Supporting Information

**Figure S1:**
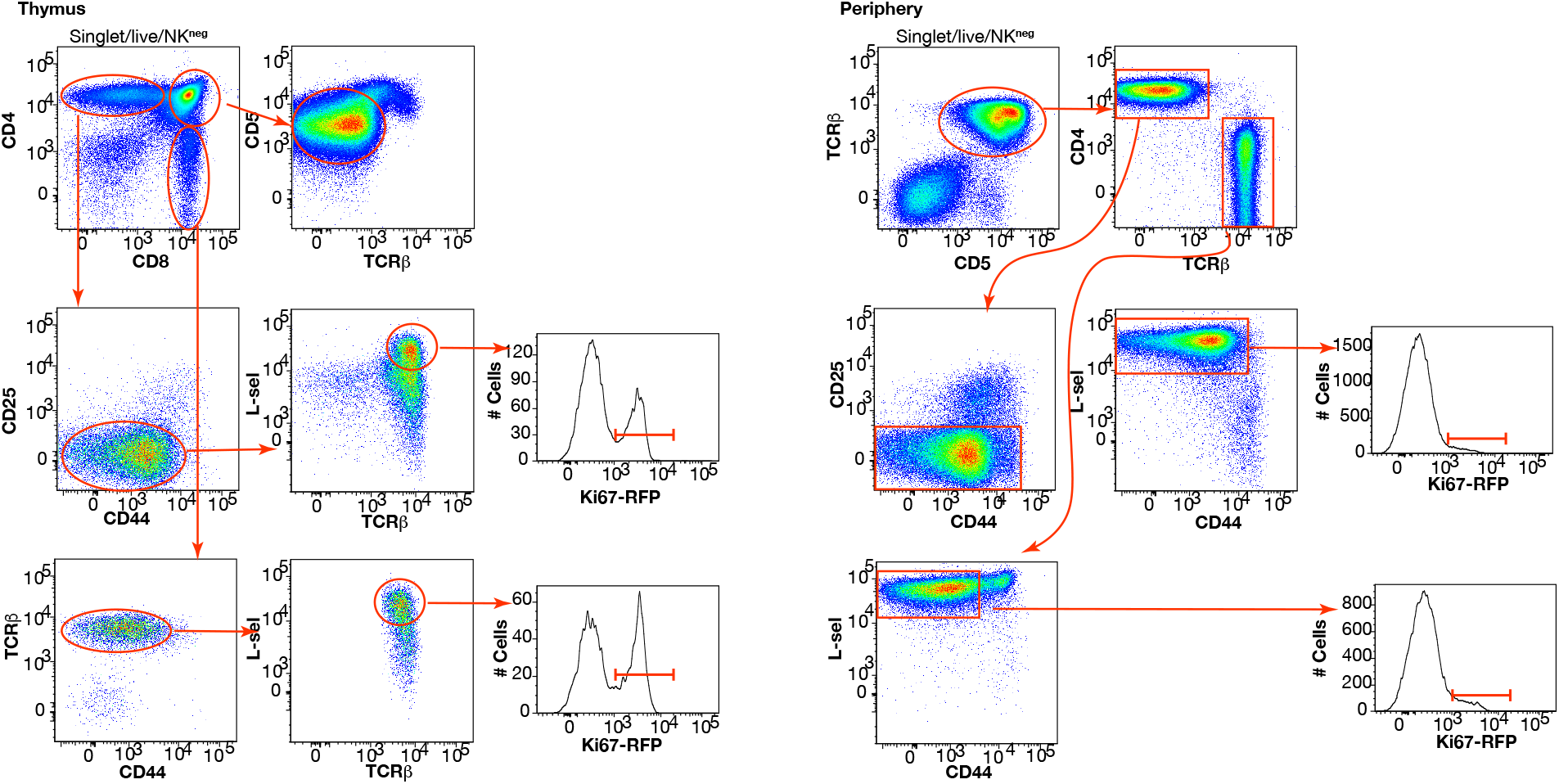
Gating strategies for thymocyte and peripheral naive T cell subsets.

**Figure S2:**
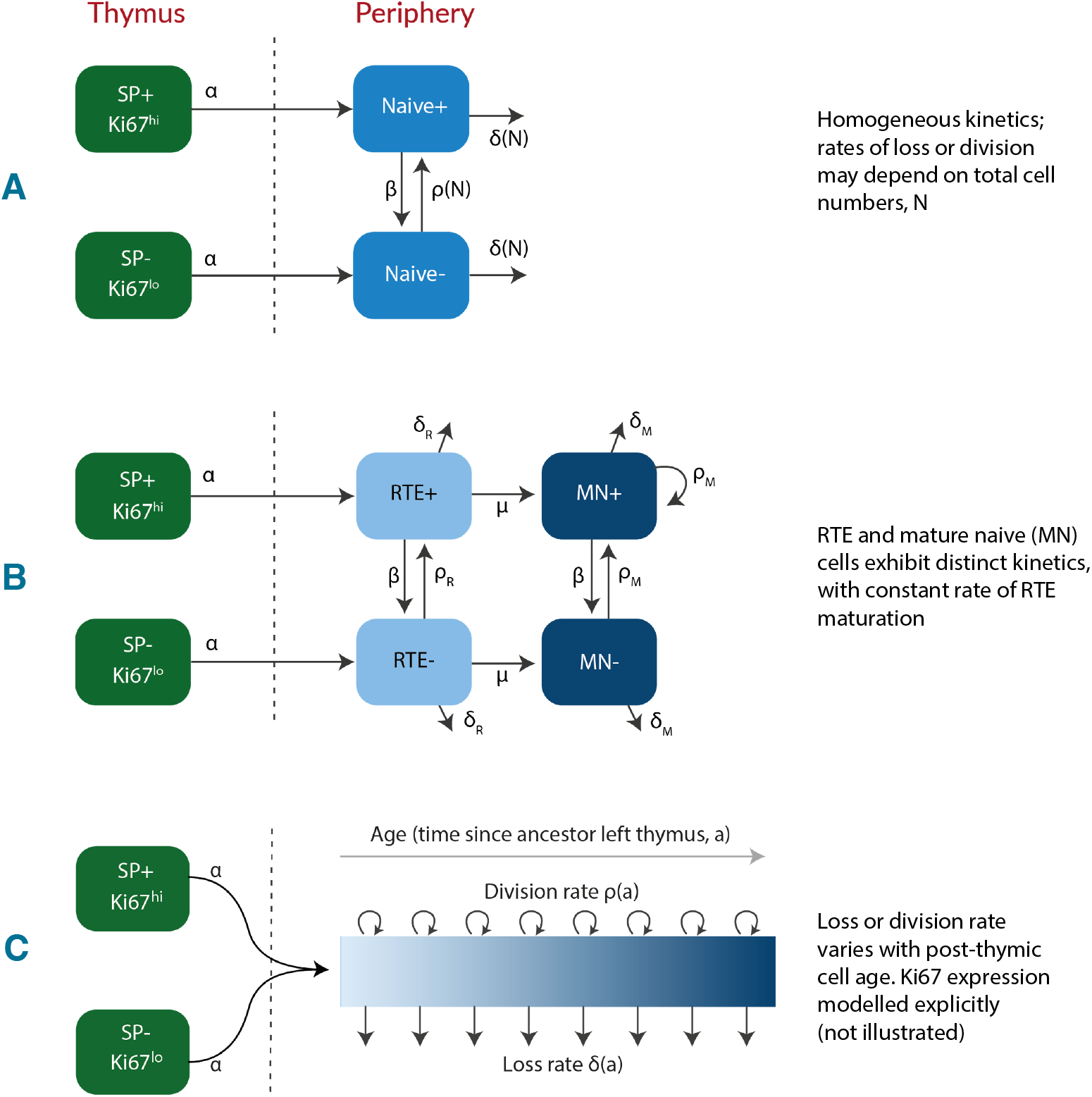
Candidate models of naive T cell dynamics. We considered three classes of model; **(A)** Kinetic homogeneity, in which all cells are lost at the same rate and divide at the same rate. In the simplest ‘neutral’ case these rates are constant. We also considered extensions in which loss or division rates were allowed to vary with total cell numbers (density-dependent models). **(B)** RTE and mature naive T cells exhibit distinct kinetics, with a constant rate of maturation. **(C)** Loss or division rates vary with post-thymic cell age. In all models we assume Ki67^low^ and Ki67^hi^ cells are exported from the thymus at rates proportional to the numbers of Ki67^low^ and Ki67^hi^ single positive thymocytes.

## Text S1 Models of naive T cell maintenance

### Text S1.1 Neutral model

We assume that the naive T cell pool is a kinetically homogeneous population that self-renews through a constant rate of division *ρ*, and is lost by death and differentiation at a constant rate *δ*. The inverse of *ρ* is the mean interdivision time, and the inverse of *δ* is the mean residence time of a cell. Thymic influx is the product of the *per capita* rate of influx *α* and the timecourse of the size of SP population *S*(*t*), which is described empirically (see Text S2). We model the dynamics of Ki67^hi^ (*N*^+^) and Ki67^low^ (*N*^−^) cells using the following ODE model;

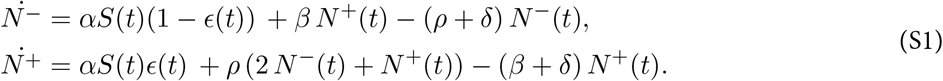

Here, *β* is the rate of loss of Ki67 expression after mitosis, and *ϵ* is the Ki67^hi^ fraction among cells immediately after export from the thymus. We assumed eqns. S1 hold identically for host and donor cells and solved them to derive the solutions to the following:

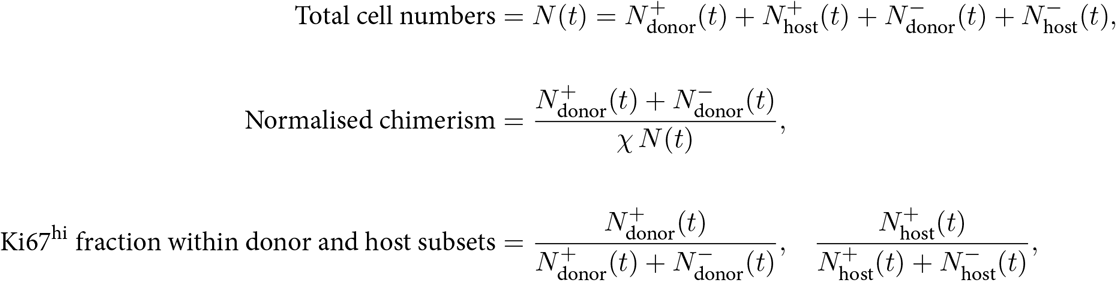

using the empirical descriptions of the size (*S*(*t*)) and Ki67^hi^ fraction (*ε*(*t*)) of SP thymocytes, their direct precursors.

### Text S1.2 Density-dependent models

In these extensions of the neutral model, the rate of cell division *ρ*, or the rate of loss *δ* vary with the size of the naive T cell population. We explored models exhibiting density-dependence in either *ρ* or *δ*. Both models assume that all cells in the population follow the same rules of self-renewal and turnover, at any given time. We defined the density-dependence using Hill functions;

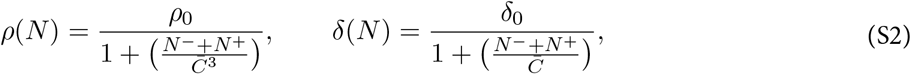

where 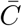 and *p*_0_ or *δ*_0_ were estimated from the model fits to the data.

### Text S1.3 RTE model

Here we treated RTE and mature naive (MN) T cells separately, allowing them to have distinct rates of division (*ρ_R_* and *ρ_N_*) and of loss (*δ_R_* and *δ_N_*). We assume a constant rate of maturation of RTE (*μ*). In this model, the expected residence time of cells in the RTE compartment is 1/(*δ_R_* + *μ*), and a proportion *μ*/(*δ_R_* + *μ*) of RTE survive to maturity. We solve the following equations for Ki67^hi^ and Ki67^low^ cells with the RTE and MN compartments, which are identical for host- and donor-derived cells;

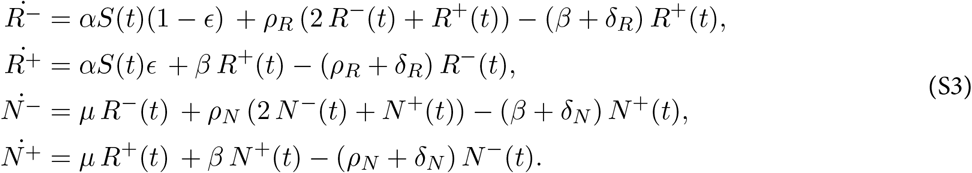

### Text S1.4 Age-dependent division and loss models

We aimed to model the population density of naive T cells *u*(*a*, *k*, *t*), where *t* is mouse age, *a* is a cell’s age (defined as the time since it or its ancestor left the thymus), and *k* is a cell’s level of Ki67 expression. We assume Ki67 expression level reaches a maximal level of *k* = 1 immediately after cell division and decays exponentially at rate *β*. We assume that the *per capita* rates at which cells divide (*ρ*) and are lost (*δ*) can be functions of mouse age *t* and/or of cell age *a*. To model the evolution of *u*(*a*, *k*, *t*), we extended an age-structured population PDE model described previously^3,22^ to include Ki67 expression.

#### Initial conditions

We assume that at some host age *t*_0_, the population has size *N*_0_ and has a cell-age distribution *γ*(*a*), where 0 ≤ *a* ≤ *t*_0_ and 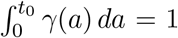 (we assume all cells are of age *a* = 0 when they leave the thymus); and these cells have a distribution of levels of Ki67 expression *ψ*(*k*), where 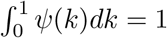 and *ψ*(*k*) = 0 for *k* ∉ (0, 1). Here for simplicity we are assuming no relation between Ki67 expression and cell age within the cells present at *t* = *t*_0_, but one could easily extend this framework with a more general initial joint distribution *P*(*a*, *k*). At times *t* ≥ *t*_0_, we assume that cells of age zero enter the naive pool from the thymus at rate *θ*(*t*) and with Ki67 distribution *φ*(*k*, *t*), where 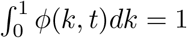 for all *t*.

#### Breaking the solution into cohorts of cells

Our approach is to track separately the fates of cells that were present at mouse age *t*_0_, who will all have age *a* > *t* – *t*_0_ at some later time t; and the fates of those that were exported from the thymus at time *t*_0_ or later, which will all have age *a* < *t* – *t*_0_. We then add these to get the full population density *u*(*a*, *k*, *t*). The master PDE for both populations combined is

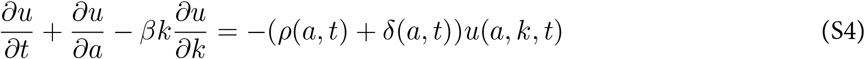

with boundary conditions

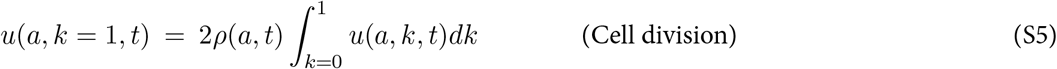

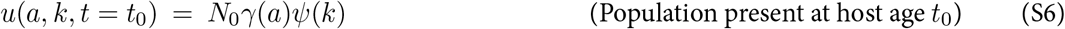

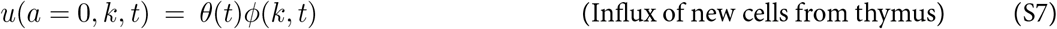

The first condition above derives from cell division; at any host age *t*, cells of age *a* at time *t* divide at rate *ρ*(*a*, *t*) generating two cells of age *a* with *k* = 1.

### Initial cohort

#### Non-divided cells

First consider those cells present at *t* = *t*_0_ that have yet to divide; this population decreases in size with a *per capita* rate *δ*(*a*, *t*) + *ρ*(*a*, *t*), and follows eqn. S4 with the single boundary condition *u*(*a*, *k*, *t* = *t*_0_) = *N*_0*γ*_(*a*)*ψ*(*k*). We solve this using the method of characteristics by identifying a variable *s* such that

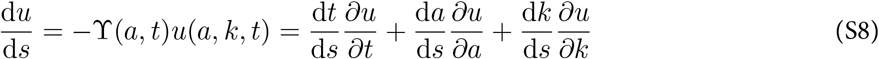

where for brevity we define *Υ*(*a*, *t*) = *ρ*(*a*, *t*) + *δ*(*a*, *t*). Equating terms with eqn. S4 gives

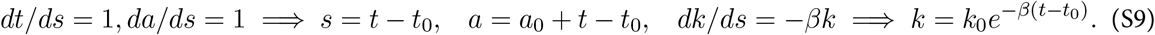

Along the characteristic that starts at (*a*_0_, *k*_0_, *t*_0_) (illustrated in Figure S3A), the population density evolves as

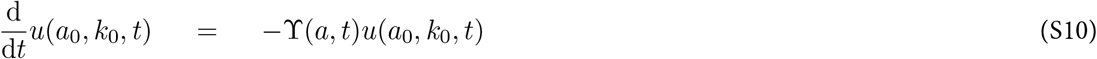

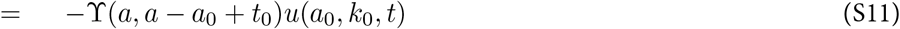

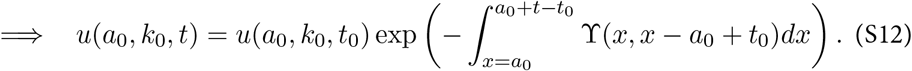

We know from eqn. S9 that *k*_0_ = *ke*^*β*(*t*–*t*_0_)^, so

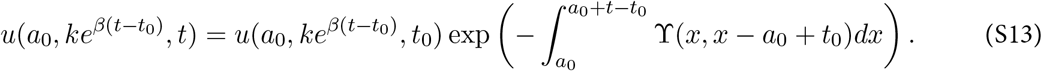

**Figure S3:**
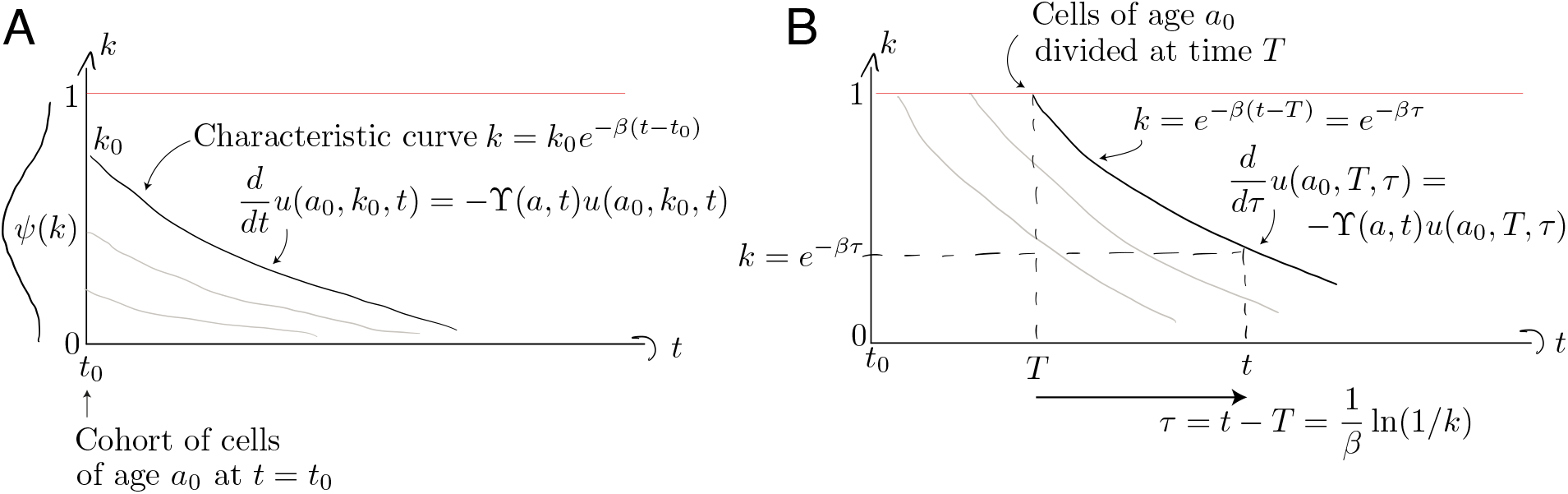
Characteristic curves for the population of age *a*_0_ present at *t* = *t*_0_ (panel A) and the population who divided at time *T* when they were of age *a*_0_ (panel B).

The population density *u*(.) must then be transformed with a Jacobian to express it as a density over *k* rather than over *ke*^*β*(*t*–*t*_0_)^. If *y* = *ke*^*β*(*t*–*t*_0_)^ then

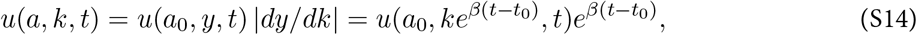

giving

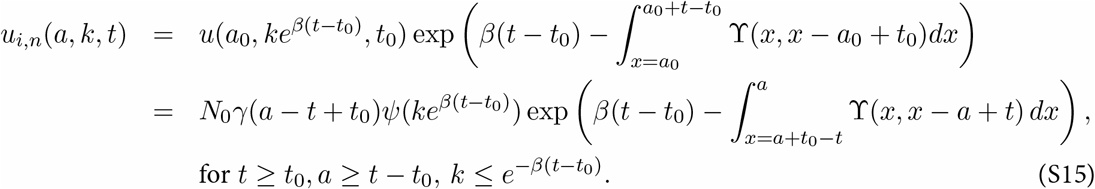

where the subscript (*i*, *n*) denotes ‘initial and non-divided’. Equation S15 holds for *k* ≤ *e*^−*β*(*t*–*t*_0_)^ because none of these cells have divided since time *t*_0_ and their Ki67 expression is decaying.

#### Divided cells

Figure S3B illustrates the evolution of cells from the initial cohort who do subsequently divide, each time resetting their Ki67 expression to *k* = 1. To follow these cohorts we solve eqn. S4 with the boundary condition describing the division of cells of age *a* at host age *t*;

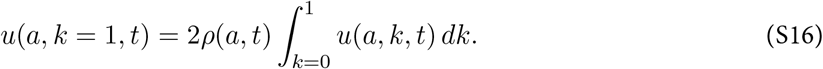

Characteristic curves are again of the general form *k* = *k*_0_*e*^−*βt*^, but we parameterise them differently to those described above, since these originate at *k* = 1 and not *t* = *t*_0_ (Figure S3B). Cells with Ki67 expression *k* must have divided a time *τ* = –(1/*β*) ln(*k*) in the past. A convenient parameterisation is therefore

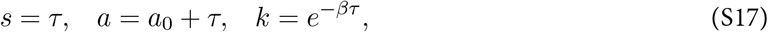

where the cells currently of age *a* divided at time *T* = *t* – *τ* when they were aged *a*_0_ = *a* – *τ*. Along these curves,

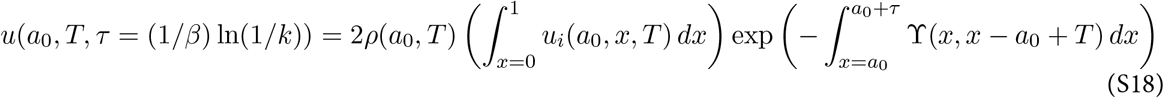

where the exponential term represents the proportion of cells on this characteristic curve that divide again or die. It integrates a cell’s experience from age *a*_0_ at host age *T*, to age *a* at host age *t* = *T* + *τ*. Here *u_i_* denotes the entire initial cohort, divided or undivided, integrated over all levels of Ki67 expression, whose cells of age *a*_0_ fed the divided population at time *T*.

To convert this to a density *u_i,d_*(*a*, *k*, *t*), we use the Jacobian *dτ*/*dk* = 1/(*βk*). This gives

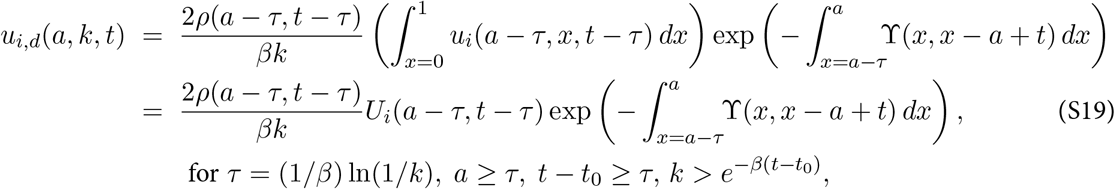

where *U_i_*(*a*, *t*) is the population density of the initial cohort (divided or undivided) at age *a* and time *t*, integrated over all *k*, which we will return to below.

### Cells that enter after *t* = *t*_0_

A similar set of calculations applies for cells that subsequently enter the pool, at which point we define them to be age zero. Again we partition these cells into those that don’t divide and those that do. For the former, consider those that are of age *a* ≤ *t* – *t*_0_ at time *t*; that is, cells that entered later than *t*_0_. We denote this population *u_θ,n_*(*a*, *k*, *t*) where *k* ≤ *e^−βa^*. These cells are what remains of the cohort exported at time *t* – *a*, at age *a* = 0, of size *θ*(*t* – *a*) and with Ki67 distribution *φ*(*ke^βa^*, *t* – *a*). These are then lost to death or division. By analogy with equation S15, the Jacobian is *e^βa^* and these non-divided cells evolve as

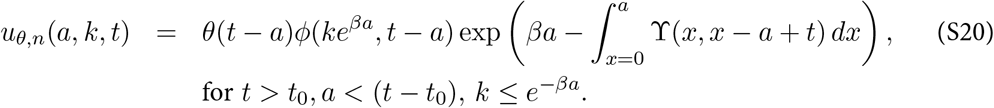

Here, cells are born with age zero and so the characteristics are *k* = *k*_0_*e^−βa^*. Similarly for divided cells; by analogy with equation S19,

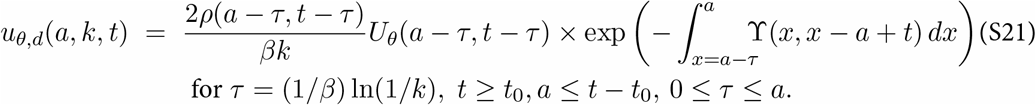

Here *U_θ_*(*a*, *t*) is the population density of cells of of age *a* at time *t*, who entered the pool a time *τ* = *t* – *a* ago and may have divided or not.

### Solving the age-structured PDE only

To complete these solutions we need the population densities *U_i_*(*a*, *t*) and *U_θ_*(*a*, *t*), ignoring Ki67 expression. These are straightforward;

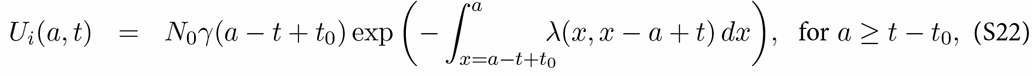

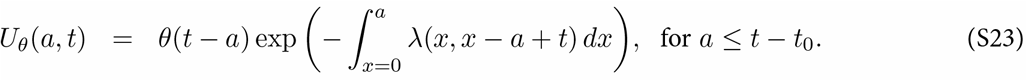

where for brevity *λ*(*a*, *t*) is the net loss rate of cells of age *a* at time *t*, which is *δ*(*a*, *t*) – *ρ*(*a*, *t*). The integral in equation S22 follows a cell whose age runs from *a* – (*t* – *t*_0_) to *a*, during which host age runs from *t* – *a* to *t*. The integral in equation S23 follows a cell whose age runs from 0 to *a*, between host ages of *t* – *a* to *t*. The two solutions join at *a* = *t* – *t*_0_; the influx at time *t*_0_ must be the density of cells of age zero in the initial cohort; *θ*(*t*_0_) = *N*_0*γ*_(0).

Therefore, to obtain the total solution *u*(*a*, *k*, *t*) = *u_i,n_* + *u_i,d_* + *u_θ,n_* + *u_θ,d_*, we add equations S15, S19, S20, S21, using the solutions for the age-structured model given in equations S22 and S23.

We can connect this solution to gated flow cytometry data by partitioning the popuation into high and low Ki67 expression. We define Ki67^hi^ cells to be those which divided no more than a time 1/*β* ago, which corresponds to a cut-off of *k* = 1/*e*. Therefore, the numbers of Ki67 positive and negative cells at time *t* are

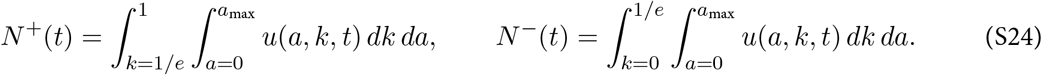

## Text S2 Constructing empirical descriptions of the dynamics of mature SP thymocytes over the mouse lifespan

We assumed that the rate of export of new naive T cells from the thymus is proportional to the numbers of single positive (SP4 and SP8) thymocytes^19^. The total numbers of SP thymocytes increase rapidly up to 6-7 weeks of age and then drop gradually over time. We used the following empirical descriptor function to capture the dynamics of thymic SP cells varying with mouse age;

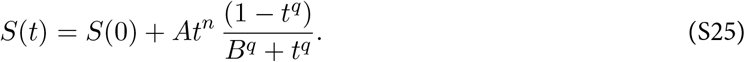

We estimated the parameters *S*(0), *A*, *n* and *q* by fitting equation S25 to the log-transformed data from wildtype mice bred in the same facility as the busulfan chimeras (Figure S4A). The rate of thymic export is then *θ*(*t*) = *αS*(*t*), with the constant *α* estimated when fitting models to the busulfan chimera data.

We modelled the Ki67^hi^ fraction within SP4 and SP8 thymocytes using the form

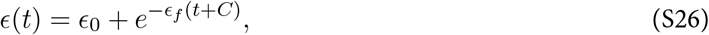

and estimated *ϵ*_0_, *ϵ_f_* and *C* by fitting this function to the logit-transformed proportions of cells that were Ki67^hi^ (Figure S4B).

When modelling data from the busulfan chimeras, we assumed that the total output from the thymus at any time is identical to that in age-matched wild type mice, but is split between donor and host cells according to the chimerism *χ* at the DP1 stage of thymic development:

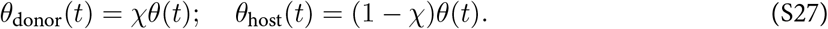

**Figure S4:**
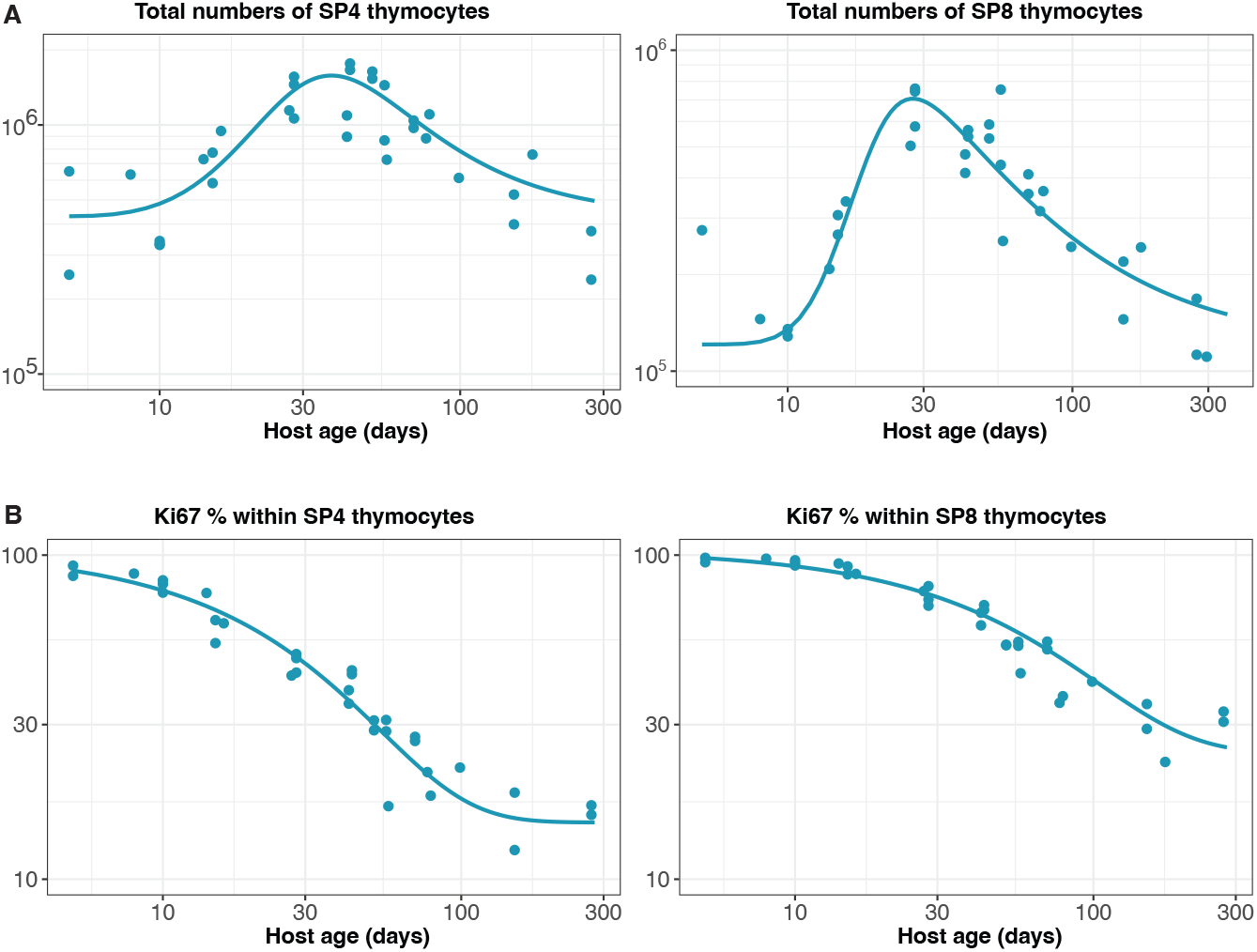
Empirical descriptions of the dynamics of the numbers and Ki67 expression of late-stage thymocytes. These curves (defined in Text S2) were used as inputs to models of the data from adult busulfan chimeric mice.

## Text S3 Model ranking and selection criteria

Each of the models described in Text S1 was fitted simultaneously to four sets of observations – cell counts, normalised chimerism, and the proportions of cells expressing Ki67 within donor and host naive T cells. To normalise residuals, cell counts were log transformed; and normalised chimerism and Ki67^hi^ fractions were logit- and arcsine square root-transformed, respectively. We used a Bayesian inference approach to estimate the model parameters and the errors associated with the measurements in each dataset. Model definitions, the prior distribution of parameters and the likelihood definitions were encoded in the *Stan* language and are available at this [Github repository](https://github.com/sanketrane/T_cell_dynamics_birth-death), and models were fitted using the Hamiltonian Monte Carlo algorithm^50^.

We compared the support for models using the leave-one-out (LOO) cross validation method. The LOO information criterion (LOO-IC) is calculated for each model by converting the expected log point-wise predictive density (*elpd*) to the deviance scale as, −2 × *elpd*^49^. The elpd estimate is a measure of the out-of-sample prediction accuracy of the model and is calculated using the *loo-2.0* package^51^ in the *Rstan* library, which employs Pareto smoothed importance sampling (PSIS)^48^ to approximate LOO cross validation. The input for this process is an array of joint likelihoods evaluated at draws from the posterior parameter distributions. ΔLOO-IC is defined as the difference in LOO-IC values between the model under consideration and the model with the lowest LOO-IC value. We then used the estimates of ΔLOO-IC to assess the relative support for models using the Pseudo-Bayesian model averaging (BMA) method^52^, implemented in the *loo-2.0* package.

**Table S1:** Ranking of models describing naive CD4 and CD8 T cell dynamics in adult busulfan chimeric mice. We fitted different instances of the three classes of model (A, B, C; illustrated in Figure S2) to data from adult busulfan chimeric mice. Measures of support relative to the best fitting model are expressed as differences in the leave-one-out information criterion (LOO-IC)^48,49^. Model weights are calculated on the basis that the relative support for two models is exp(−ΔLOO-IC); so, for instance, a difference in LOO-IC of 6 or more implies at least 20-fold lower support for the model with the larger LOO-IC value.

| Population | Model | $\Delta$ LOO-IC | Model weight (%) |
| --- | --- | --- | --- |
| Naive CD4 | C – Loss varying with cell age | 0 | 86.3 |
|  | C – Division varying with cell age | 21 | 13 |
|  | A – Neutral | 38 | 0.5 |
|  | B – Distinct dynamics of RTE mature naive T cells | 40 | 0.2 |
|  | A – Density dependent loss | 48 | 0.0 |
|  | A – Density dependent division (LIP) | 58 | 0.0 |
| Naive CD8 | C – Division varying with cell age | 0 | 85.0 |
|  | C – Loss varying with cell age | 13 | 9.0 |
|  | A – Density dependent division (LIP) | 19 | 4.5 |
|  | A – Density dependent loss | 25 | 1.5 |
|  | B – Distinct dynamics of RTE mature naive T cells | 26 | 0.0 |
|  | A – Neutral | 27 | 0.0 |

**Figure S5:**
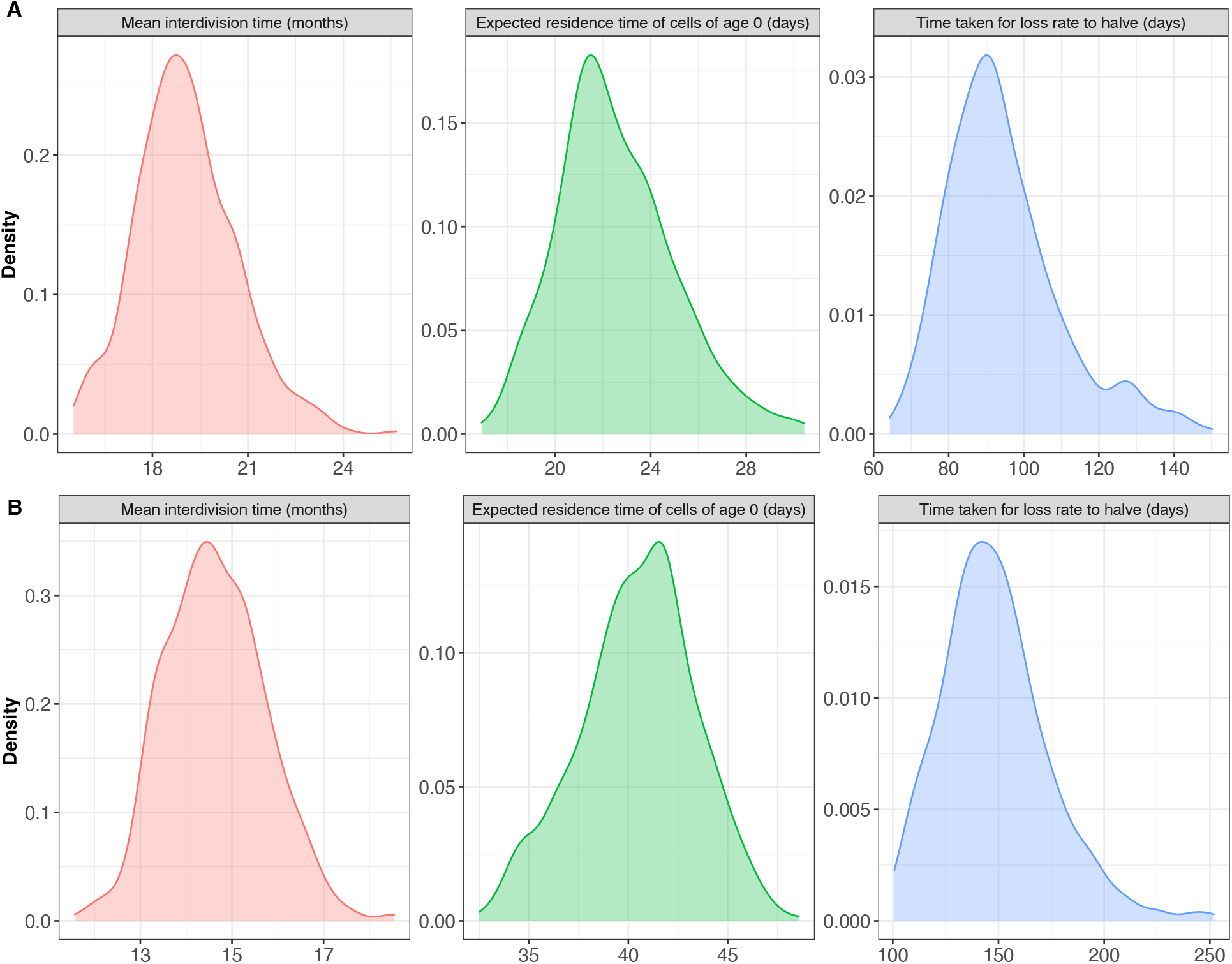
Posterior distributions of key parameters for naive **(A)** CD4 and **(B)** CD8 T cells derived from fitting the age-dependent loss model to the data from adult busulfan chimeric mice.

## Text S4 Extending models back to near-birth to predict the dynamics of naive T cells in neonates

In order to model the busulfan chimera data, in which mice underwent BMT at different ages, we needed to define the healthy dynamics of naive T cells. We took the approach of evolving the naive T cell pool from 1 day of age assuming that kinetic parameters were constant across the lifespan. In this way, every mouse began from the same ‘baseline’ state just after birth, and varied only in its age at BMT and in the level of stable chimerism within the bone marrow and thymus. The free parameters for this aspect of the model were then the (unknown) numbers of Ki67^hi^ and Ki67^low^ naive CD4 or CD8 T cells in a 1-day-old mouse, 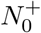 and 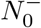 respectively. For each model, we then calculated the predicted numbers of host-derived Ki67^hi^ and Ki67^low^ cells at the age of BMT (*t*_BMT_) by simulating forward from this initial condition, allowing for the continued influx of Ki67^hi^ and Ki67^low^ cells from the thymus as described in Text S2.

In cell-age dependent models, we also defined the Ki67 distribution within the pre-existing naive T cells (initial cohort) and among cells subsequently exported from the thymus emigrants. We also assumed a uniform distribution of cell ages (*γ*(*a*) = 1) in 1 day old mice.

### Initial cohort

Naive T cells present at mouse age *t*_0_ = 1 day are enriched with Ki67^hi^ cells (Figure 4A and B), and we defined the distribution of their normalized Ki67 expression *k* over (0,1] to be *ψ*(*k*) = ^*ek*^/(*e^k^*–1). We assume Ki67 is lost exponentially at a constant rate *β*, such that at a time *s* after division *k*(*s*) = *e^−βs^*. Our results were not sensitive to the form of *ψ*(*k*) since it ‘washes out’ on a timescale of 1/*β* = 3.5 days.

We define the cut-off that separates Ki67^hi^ from Ki67^low^ cells to be 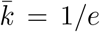, such that cells spend a time 1/*β* as Ki67^hi^. The fraction of Ki67^hi^ cells in the initial cohort is then obtained by integrating the initial Ki67 distribution *ψ*(*k*) from 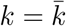 to *k* = 1, yielding

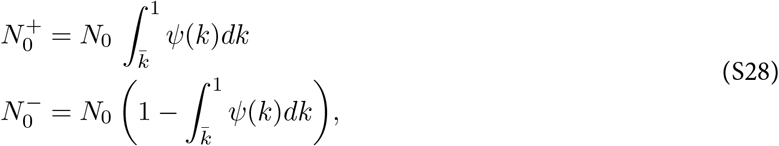

where *N*_0_ is the total number of naive CD4 or CD8 T cells present in a 1-day old mouse, and was a free parameter in the models.

### Naive T cells that subsequently enter the periphery

The Ki67^hi^ fraction within SP thymocytes (*ϵ*(*t*); equation S26) varies with time, which implies that the distribution of Ki67 expression within naive T cells of age zero (*i.e*., just exported from the thymus) also change with time. We define this distribution to be *φ*(*k*, *t*),such that the Ki67^hi^ fraction among new RTE at mouse age *t* is

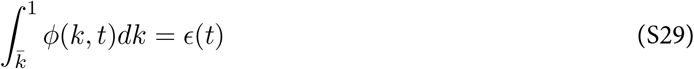

We then defined an empirical step function *φ*(*k*, *t*) to be consistent with this relationship;

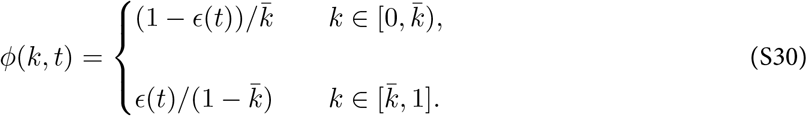

To generate the predicted numbers and Ki67^hi^ fractions of naive CD4 and CD8 T cells from age 5 days onwards (Figure 4), it was then straightforward to plot the model fits derived from the adult busulfan chimeric mice; note that none of the data derived from the younger, wild-type mice were used in the fitting.

## Text S5 Predicting RTE dynamics in Rag/Ki67 dual reporter mice

In Rag^GFP^ Ki67^RFP^ reporter mice, RTE are identified based on the transient expression of GFP, which we assume decays with first order kinetics. Our models estimated very low rates of division among naive CD4 and CD8 T cells, such that any dilution of GFP through division is minimal. There is therefore a simple and direct correlation between GFP expression *f* and cell age *a* within naive T cells, which we used to predict the fractions of GFP^+^ cells (*F*) within the naive compartment.

Using the favoured age-structured models described in Text S1.4, the age distribution of the naive T cell pool at mouse age *t* is given by equations S22 and S23 as *U*(*a*, *t*) = *U_i_*(*a*, *t*) + *U_θ_*(*a*, *t*). The fraction of cells that are GFP^+^ is then

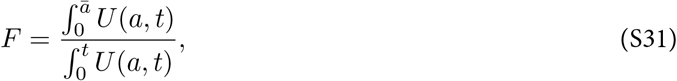

where *ā* is the (unknown) time required for cells to transition from GFP^+^ to GFP^−^. Implicit in this calculation is the assumption that GFP levels are similar in Ki67^low^ and Ki67^hi^ RTE; mature SP cells are indeed very bright for GFP with minimal differences when stratified by Ki67 expression (data not shown). We then used the parameters derived from busulfan chimera data to generate *U*(*a*, *t*), and estimated *ā* by fitting equation S31 to the timecourse of the GFP^+^ fraction within naive T cells observed in Rag/Ki67 dual reporter mice. The fits are shown in Figure 4C and D, red lines in the leftmost panels. We then generated the predicted timecourses of the GFP^+^ Ki67^+^ and GFP^+^ Ki67^+^ fractions by multiplying *F* with the predictions of Ki67^hi^ and Ki67^low^ fractions that we derived by running the age-dependent loss model from age of the youngest GFP/Ki67 reporter mouse (11 days).

## Text S6 Hierarchical modelling of naive CD8 T cell timestamping data

The data from Reynaldi *et al*.^23^, shown in Figure 5, comprised longitudinal samples drawn from animals in five different age groups who were each treated with pulses of Tamoxifen to label cohorts of CD8 T cells leaving the thymus. To use these data to estimate how cell loss rates vary as a function of both cell and host age, we took a hierarchical modelling approach. We allowed for animal and/or group-level variation in the initial numbers of RFP labelled cells in each animal (*N*_0_) and in their initial loss rate (that is, the instantaneous net loss rate of cells of age zero, just exported from the thymus). We began by modeling the kinetics of labelled cells with the assumption that their net loss rate *λ* varies with their post-thymic age *a* as *λ*(*a*) = *λ*_0*e*^−*γa*^_. The population density of cells of age *a* at mouse age *t*, *N*(*a*, *t*), then obeys

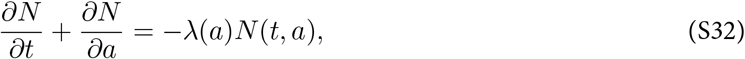

with the boundary condition *N*(*T*, *a*) = *N*_0_*δ*(*a*), where *T* is the mouse age at the time of treatment and *δ*(.)is the Dirac delta function. For mouse *i*, in age group *j*, at timepoint *k*, the observed cell numbers are *y_ijk_*, where

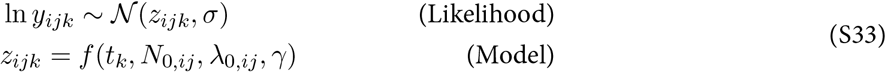

We considered models in which the initial cell numbers *N*_0_ and initial loss rate *λ*_0_ for each mouse were either drawn from a single parent distribution or from distributions with means and variances specific to each age group. We defined priors for *μ_N_*, *σ_N_*, *μ_λ_*, *σ_λ_* and *γ* and fitted permutations of the hierarchical age-structured model to the time courses of labelled CD8 T cell numbers (Table S34). The best-fitting model exhibited group-specific values of the initial RTE loss rate *λ*_0_, and variation in the initial numbers of labelled cells (*N*_0_) across mice, likely deriving from variations in the efficiency of tamoxifen-driven labelling.

**Table S2:** Comparing support for hierarchical age-structured models of the data from ref. 23.

| Model | Initial numbers | Net loss rate at age 0 | $\Delta$ LOO-IC | Weight % |
| --- | --- | --- | --- | --- |
| $N_0$ varying at animal level | $N_{0,i} \sim \mathcal{N}(\mu_N, \sigma_N)$ | constant | 316 | 0.0 |
| $N_0$ varying at animal level; $\lambda_0$ varying at group level | $N_{0,i} \sim \mathcal{N}(\mu_N, \sigma_N)$ | $\lambda_{0,j} \sim \mathcal{N}(\mu_\lambda, \sigma_\lambda)$ | 0.0 | 100 |
| $N_0$ varying at animal level; $\lambda_0$ varying at animal level | $N_{0,i} \sim \mathcal{N}(\mu_N, \sigma_N)$ | $\lambda_{0,i} \sim \mathcal{N}(\mu_\lambda, \sigma_\lambda)$ | 73 | 0.0 |
| $N_0$ varying at group level; $\lambda_0$ varying at animal level | $N_{0,j} \sim \mathcal{N}(\mu_N, \sigma_N)$ | $\lambda_{0,i} \sim \mathcal{N}(\mu_\lambda, \sigma_\lambda)$ | 313 | 0.0 |

We then generated an explicit, empirical description of the variation in *λ*_0_ with mouse age t, using the group-specific estimates of *λ*_0_ from the best-fitting hierarchical model. We used the following model of the net loss rate of cells of age *a* at time *t*, with estimated parameters *λ_h_*,*Q*,*q* and *γ*;

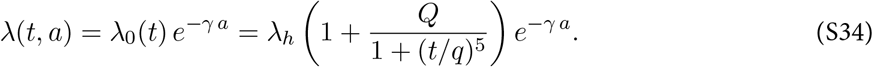

**Figure S6:**
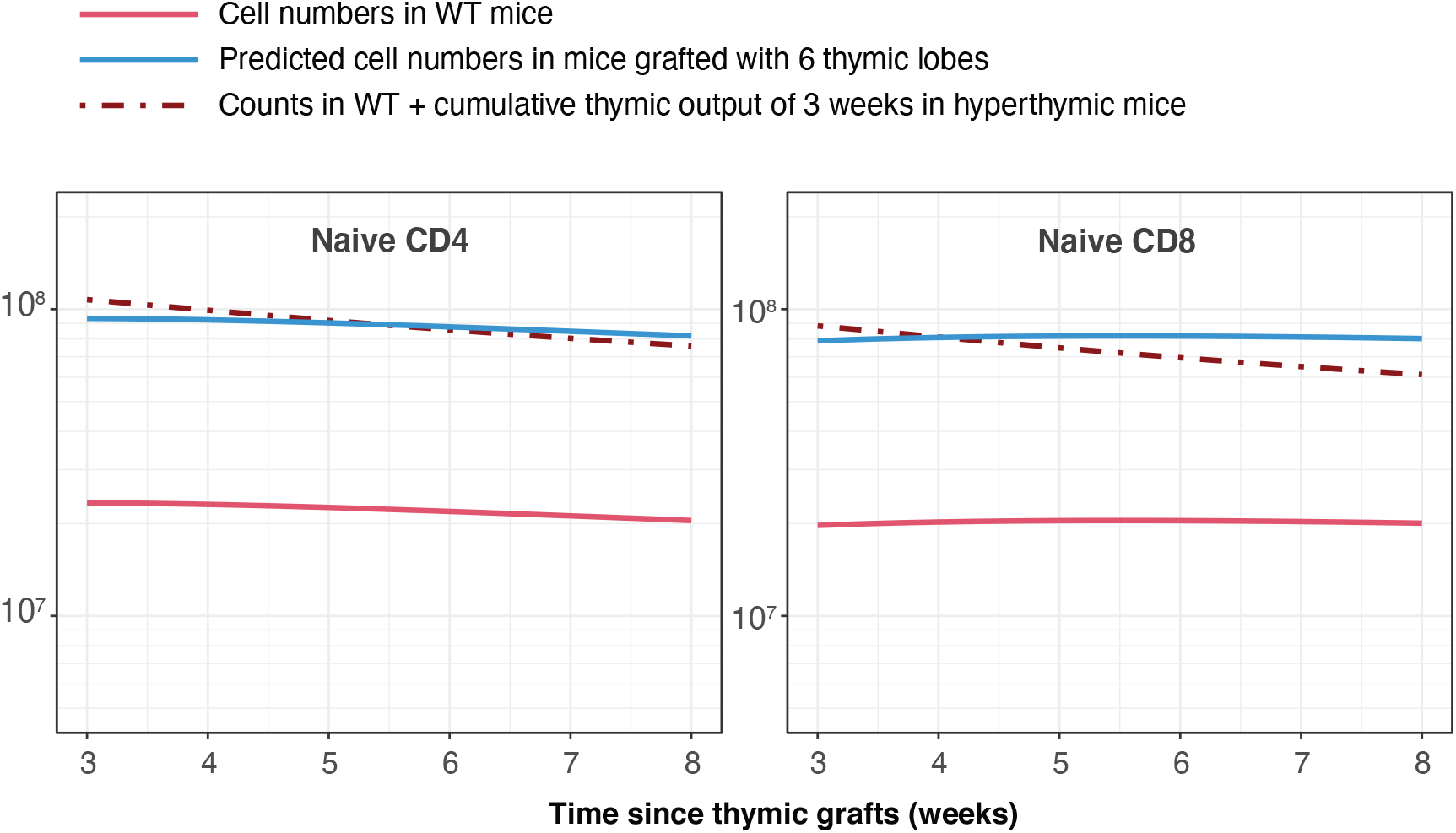
Simulating the outcome of transplanting 6 additional thymi, as described by Berzins *at al*.^20^. The change in numbers of naive CD4 and CD8 T cells is equivalent to 3 weeks of thymic output.

## References

1. Scollay RG, Butcher EC, Weissman IL (1980) Thymus cell migration. Quantitative aspects of cellular traffic from the thymus to the periphery in mice. Eur J Immunol 10(3):210–8.

2. den Braber I, et al. (2012) Maintenance of peripheral naive T cells is sustained by thymus output in mice but not humans. Immunity 36(2):288–97.

3. Hogan T, Gossel G, Yates AJ, Seddon B (2015) Temporal fate mapping reveals age-linked heterogeneity in naive T lymphocytes in mice. Proc Natl Acad Sci U S A 112(50):E6917–26.

4. Egerton M, Scollay R, Shortman K (1990) Kinetics of mature T-cell development in the thymus. Proc Natl Acad Sci U S A 87(7):2579–82.

5. Graziano M, St-Pierre Y, Beauchemin C, Desrosiers M, Potworowski EF (1998) The fate of thymocytes labeled in vivo with CFSE. Exp Cell Res 240(1):75–85.

6. Rocha B, Dautigny N, Pereira P (1989) Peripheral T lymphocytes: expansion potential and homeostatic regulation of pool sizes and CD4/CD8 ratios in vivo. Eur J Immunol 19(5):905–11.

7. Almeida AR, Borghans JA, Freitas AA (2001) T cell homeostasis: thymus regeneration and peripheral T cell restoration in mice with a reduced fraction of competent precursors. J Exp Med 194(5):591–9.

8. Yates A, Saini M, Mathiot A, Seddon B (2008) Mathematical modeling reveals the biological program regulating lymphopenia-induced proliferation. J Immunol 180(3):1414–1422.

9. Houston, Jr EG, Higdon LE, Fink PJ (2011) Recent thymic emigrants are preferentially incorporated only into the depleted T-cell pool. Proc Natl Acad Sci U S A 108(13):5366–71.

10. Hogan T, et al. (2013) Clonally diverse T cell homeostasis is maintained by a common program of cellcycle control. J Immunol 190(8):3985–93.

11. Min B, et al. (2003) Neonates support lymphopenia-induced proliferation. Immunity 18(1):131–40.

12. Le Campion A, et al. (2002) Naive T cells proliferate strongly in neonatal mice in response to self-peptide/self-MHC complexes. Proceedings of the National Academy of Sciences 99(7):4538–4543.

13. Dzierzak E, Daly B, Fraser P, Larsson L, Müller A (1993) Thy-1 tk transgenic mice with a conditional lymphocyte deficiency. Int Immunol 5(8):975–84.

14. Hogan T, et al. (2019) Differential impact of self and environmental antigens on the ontogeny and maintenance of CD4+ T cell memory. eLife 8.

15. Akue AD, Lee JY, Jameson SC (2012) Derivation and maintenance of virtual memory CD8 T cells. J Immunol 188(6):2516–23.

16. Houston, Jr EG, Nechanitzky R, Fink PJ (2008) Cutting edge: Contact with secondary lymphoid organs drives postthymic T cell maturation. J Immunol 181(8):5213–7.

17. Adkins B (1999) T-cell function in newborn mice and humans. Immunol Today 20(7):330–5.

18. Wang J, et al. (2016) Fetal and adult progenitors give rise to unique populations of CD8+ T cells. Blood 128(26):3073–3082.

19. Berzins SP, Boyd RL, Miller JF (1998) The role of the thymus and recent thymic migrants in the maintenance of the adult peripheral lymphocyte pool. J Exp Med 187(11):1839–48.

20. Berzins SP, Godfrey DI, Miller JF, Boyd RL (1999) A central role for thymic emigrants in peripheral T cell homeostasis. Proc Natl Acad Sci U S A 96(17):9787–91.

21. van Hoeven V, et al. (2017) Dynamics of Recent Thymic Emigrants in Young Adult Mice. Front Immunol 8:933.

22. Rane S, Hogan T, Seddon B, Yates AJ (2018) Age is not just a number: Naive T cells increase their ability to persist in the circulation over time. PLoS Biol 16(4):e2003949.

23. Reynaldi A, et al. (2019) Fate mapping reveals the age structure of the peripheral T cell compartment. Proc Natl Acad Sci U S A.

24. Johnson PLF, Yates AJ, Goronzy JJ, Antia R (2012) Peripheral selection rather than thymic involution explains sudden contraction in naive CD4 T-cell diversity with age. Proc Natl Acad Sci U S A 109(52):21432–7.

25. Mold JE, et al. (2019) Cell generation dynamics underlying naive T-cell homeostasis in adult humans. PLOS Biology 17(10):e3000383.

26. Hogan T, Yates A, Seddon B (2017) Generation of Busulfan chimeric mice for the analysis of T cell population dynamics. Bio-protocol 4(24).

27. Verheijen M, Rane S, Pearson C, Yates AJ, Seddon B (2020) Fate Mapping Quantifies the Dynamics of B Cell Development and Activation throughout Life. Cell Reports 33(7):108376.

28. Gossel G, Hogan T, Cownden D, Seddon B, Yates AJ (2017) Memory CD4 T cell subsets are kinetically heterogeneous and replenished from naive T cells at high levels. eLife 6.

29. Miller I, et al. (2018) Ki67 is a graded rather than a binary marker of proliferation versus quiescence. Cell Rep 24(5):1105–1112.e5.

30. Boursalian TE, Golob J, Soper DM, Cooper CJ, Fink PJ (2004) Continued maturation of thymic emigrants in the periphery. Nature Immunology 5(4):418 – 425.

31. Basak O, et al. (2014) Mapping early fate determination in lgr5+ crypt stem cells using a novel ki67-rfp allele. The EMBO Journal 33(18):2057–2068.

32. Modigliani Y, Coutinho G, Burlen-Defranoux O, Coutinho A, Bandeira A (1994) Differential contribution of thymic outputs and peripheral expansion in the development of peripheral T cell pools. Eur J Immunol 24(5):1223–7.

33. Gonçalves P, et al. (2017) A new mechanism shapes the naïve CD8 T cell repertoire: the selection for full diversity. Molecular Immunology 85:66–80.

34. Quigley M, et al. (2010) Convergent recombination shapes the clonotypic landscape of the naïve T-cell repertoire. Proc Natl Acad Sci U S A 107(45):19414–19419.

35. Qi Q, et al. (2014) Diversity and clonal selection in the human T-cell repertoire. Proc Natl Acad Sci U S A 111(36):13139–13144.

36. de Greef PC, et al. (2020) The naive T-cell receptor repertoire has an extremely broad distribution of clone sizes. eLife 9.

37. Mora T, Walczak AM (2019) How many different clonotypes do immune repertoires contain? Current Opinion in Systems Biology 18:104–110.

38. Tsukamoto H, Huston GE, Dibble J, Duso DK, Swain SL (2010) Bim dictates naive CD4 T cell lifespan and the development of age-associated functional defects. The Journal of Immunology 185(8):4535–4544.

39. Dowling MR, Milutinović D, Hodgkin PD (2005) Modelling cell lifespan and proliferation: is likelihood to die or to divide independent of age? J R Soc Interface 2(5):517–26.

40. Seddon B, Zamoyska R (2002) TCR signals mediated by Src family kinases are essential for the survival of naive T cells. J Immunol 169(6):2997–3005.

41. Martin B, Bécourt C, Bienvenu B, Lucas B (2006) Self-recognition is crucial for maintaining the peripheral CD4+ T-cell pool in a nonlymphopenic environment. Blood 108(1):270–7.

42. Tsukamoto H, et al. (2009) Age-associated increase in lifespan of naive CD4 T cells contributes to T-cell homeostasis but facilitates development of functional defects. Proc Natl Acad Sci U S A 106(43):18333–8.

43. Cho BK, Rao VP, Ge Q, Eisen HN, Chen J (2000) Homeostasis-stimulated proliferation drives naive T cells to differentiate directly into memory T cells. J Exp Med 192(4):549–56.

44. Min B, Paul WE (2005) Endogenous proliferation: burst-like CD4 T cell proliferation in lymphopenic settings. Semin Immunol 17(3):201–7.

45. Hazenberg MD, et al. (2000) T-cell division in human immunodeficiency virus (HIV)-1 infection is mainly due to immune activation: a longitudinal analysis in patients before and during highly active antiretroviral therapy (HAART). Blood 95(1):249–55.

46. Baliu-Piqué M, et al. (2021) Cell-density independent increased lymphocyte production and loss rates post-autologous HSCT. Elife 10.

47. Dutilh BE, de Boer RJ (2003) Decline in excision circles requires homeostatic renewal or homeostatic death of naive T cells. J Theor Biol 224(3):351–8.

48. Vehtari A, Gelman A, Gabry J (2015) Efficient implementation of leave-one-out cross-validation and WAIC for evaluating fitted Bayesian models. Statistics and Computing, arXiv:1507.04544 27(5):1413–1432.

49. Vehtari A, Gelman A, Gabry J (2016) Practical bayesian model evaluation using leave-one-out crossvalidation and WAIC. Statistics and Computing 27(5):1413–1432.

50. Stan Development Team (2022) Stan Modeling Language Users Guide and Reference Manual.

51. Vehtari A, et al. (2020) loo: Efficient leave-one-out cross-validation and WAIC for Bayesian models (R package version 2.4.1).

52. Yao Y, Vehtari A, Simpson D, Gelman A (2018) Using Stacking to Average Bayesian Predictive Distributions (with Discussion). Bayesian Analysis 13(3).

